# THERAPY RESISTANCE IN MELANOMA DRIVES CILIOGENESIS AND GLUTAMINE/ASPARTATE METABOLISM DEPENDENCY

**DOI:** 10.64898/2026.10.13.758838

**Authors:** Pietro Lugato, Raul Castro-Portuguez, Christine Francesca Thomas, Silvia Pedretti, Roxana E. Oberkersch, Nico Mitro, Massimo Mattia Santoro

## Abstract

Despite the remarkable success of new therapeutic approaches, a considerable number of melanoma patients still relapse on conventional therapies within two years and develop resistance. Alterations in metabolism, including amino acid availability and regulation of metabolic processes, have been implicated in acquired drug resistance. In this study, we developed in vitro and in vivo models of target therapy resistance (TTR) and identified a genetic program that supports primary ciliogenesis in drug-resistant BRAF melanoma. We established that drug resistance enhances ciliogenesis, which is needed to respond and survive to combination therapy (CT). Mechanistically, impairing cilia formation (IFT88^KO^) rewires glutamine and aspartate metabolism. Overall, we provide a new molecular mechanism underlying therapy resistance that should be considered to tailored therapeutic strategies.

## Introduction

Melanomas are one of the most aggressive types of tumors in humans (Arnold *et al*, 2022; Caraviello *et al*, 2025). Despite remarkable success with new forms of treatment such as immunotherapies, there are still many melanoma patients who cannot be cured or who later suffer a relapse of the disease following successful treatment (Peirano *et al*, 2023). About 50% of human melanomas carry BRAF mutations, and BRAFV600E is the most prevalent mutation, which results in constitutively overactive MAPK–MEK–ERK signaling (Alexandrov *et al*, 2020; Board *et al*, 2009). Although combined BRAF and MEK inhibition induces rapid tumor regression, durable responses are uncommon, as most patients relapse due to acquired resistance, proving that the lack of a complete response remains a major issue (Robert *et al*, 2015; Sun *et al*, 2014; Tanda *et al*, 2020). A key driver of resistance is melanoma phenotypic plasticity, which enables non-genetic adaptation through coordinated transcriptional, signaling, translational, and metabolic reprogramming (Shen *et al*, 2025; Tan *et al*, 2024). While transcriptional and signaling mechanisms have been extensively characterized, the contribution of selective mRNA translation and its interplay with especially amino acid metabolism rewiring remains poorly understood (Robertson *et al*, 2024).

Primary cilia are sensory organelles that protrude from the cell membrane. From this extroverted position, they respond to changes in the cellular environment and convey various signaling pathways to control cell function, such as cell maintenance (Mill *et al*, 2023; Reiter & Leroux, 2017). Primary cilia are present throughout the body, from kidney tubular epithelium and biliary ducts to chondrocytes, fibroblasts, neurons, and endocrine cells, reflecting their broad, evolutionarily conserved importance in tissue development and homeostasis and serving as a cell- and context-specific signaling and sensing hub. This organelle plays a wide variety of physiological roles relevant to differentiation, development, and tissue homeostasis (Oh *et al*, 2015). Cilia play a critical role in some cancers by providing a spatially localized platform for Hedgehog, Notch, WNT, and some receptor tyrosine kinase signaling, as well as mechanosensation (Collinson & Tanos, 2025; Liu *et al*, 2018). The functions of primary cilia in melanocytes and melanoma cells are, however, poorly characterized. One study demonstrated that the loss of primary cilia is strongly correlated with melanoma (and showed that the loss of cilia in melanoma is unlikely to be caused by increased proliferation and cell cycle progression) (Snedecor *et al*, 2015). A second study showed that amplification of EZH2 in melanoma leads to cilia loss that, in turn, enhances pro-tumorigenic WNT/β-catenin signaling and is itself sufficient to drive metastatic melanoma in benign cells (Zingg *et al*, 2018).

Therapeutic resistance is frequently associated with the reprogramming of metabolic pathways in cancer cells (Finley, 2023; Luís *et al*, 2020). The rapid proliferation of melanoma cells is supported by enhanced aerobic glycolysis, which is facilitated by the BRAFV600E mutation-induced activation of the BRAF/MEK/ERK pathway (Hall *et al*, 2013; Haq *et al*, 2013). Conversely, BRAFV600E inhibition shifts metabolism from aerobic glycolysis toward increased mitochondrial respiration, increasing reactive oxygen species (ROS) production and altering the redox environment (Corazao-Rozas *et al*, 2013). Reconfiguring metabolic and signaling pathways within drug-tolerant cells is currently the subject of mechanistic investigations into resistance to cancer targeted therapies. Recently, an interesting link between cilia and metabolism has been proposed in the context of glycolysis and glutamine anaplerosis (Jenks *et al*, 2018; Steidl *et al*, 2023; Teperino *et al*, 2012). Defining the functions of primary cilia in melanoma cells will undoubtedly contribute to our understanding of therapy resistance.

Here, we identify changes in primary cilia formation during resistance to BRAFi+MEKi combination therapy in melanoma. Our bulk RNA sequencing approach, combined with RiboSeq analysis, revealed specific transcriptional and translational changes in drug-resistant cells, leading to ciliogenesis. We utilize a targeted metabolomic approach to identify changes in glutamine and aspartate metabolism driven by cilia to escape cell death. Finally, we explore the possibility of combining amino acid starvation as well as defective primary cilia formation to enhance CT against drug-resistant tumors, thus unveiling a novel strategy to combat drug resistance in melanoma.

## RESULTS

### BRAFV600E melanoma tumors undergoing targeted combination therapy (CT) display globally reduced mRNA translation

To identify new mechanisms that could contribute to resistance in BRAFV600E melanoma, we analyzed melanoma RNA expression profiling data from patients who progressed on Dabrafenib (BRAFi) and Trametinib (MEKi) therapy and identified activated and suppressed transcriptional programs driving resistance to combination BRAFi plus MEKi therapy (GSE77940 dataset) (Wagle *et al*, 2014). Gene set enrichment analysis (GSEA) using KEGG and REACTOME databases highlights that “ribosome” and “eukaryotic translation” pathways are the most suppressed in patients resistant to combined BRAFi and MEKi treatment (Figures 1A and S1A). Because bulk RNA sequencing data came from tumor tissue biopsies, where both cancer and stromal cells contribute to gene expression patterns, we next tested whether the same BRAFi and MEKi combination therapy (CT: Dabrafenib + Trametinib) yields the same trend in melanoma cells alone using a BRAFV600E melanoma cell culture model. Thus, we developed a model of acquired resistance by chronically treating A375 BRAFV600E-mutated melanoma cells with the combination therapy (CT) of Dabrafenib and Trametinib. Cells were maintained in CT until they could proliferate again under continued drug pressure, after overcoming the initial slow-cycling drug-tolerant phase. We named these drug-resistant cells (A375DR) to distinguish them from naïve, non-resistant counterparts (A375N). We performed GSEA on bulk RNA sequencing data of A375DR vs A375N cells and identified “ribosome biogenesis” and “aminoacyl-tRNA biosynthesis” to be among the suppressed gene sets (KEGG) in A375DR cells (Figure 1B). Similar results were observed when using the Gene Ontology (GO) database of biological processes (GO-BP) (Figure S1B).

**Figure 1.**
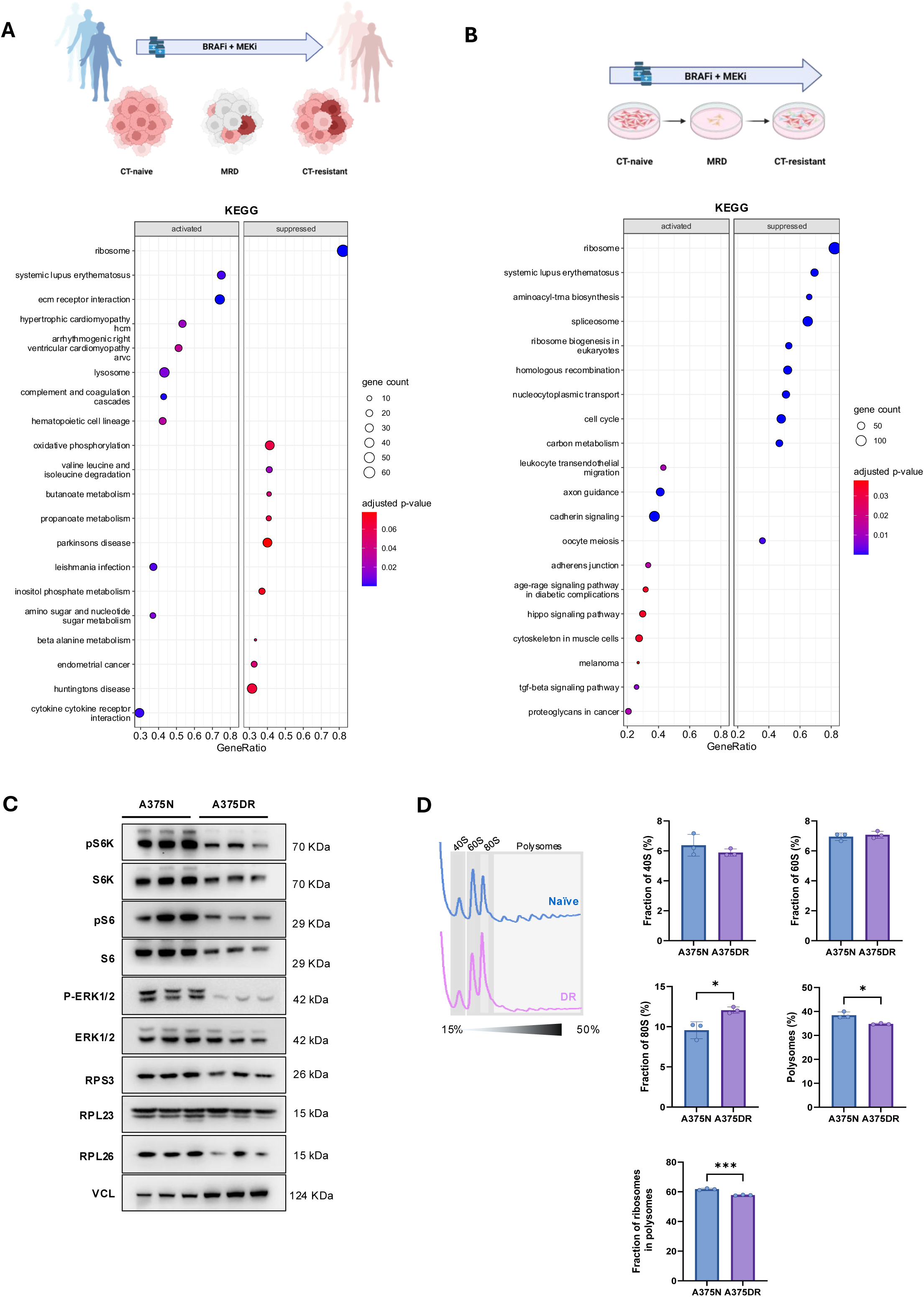
BRAF melanoma undergoing CT display globally reduced mRNA translation. A) Schematic overview of the data from GSE77940. Dot plot showing the gene sets significantly enriched in post-treatment melanoma samples compared to pre-treatment melanoma samples in the dataset GSE77940 detected by gene set enrichment analysis (GSEA) using the KEGG database. Results are split into activated and suppressed processes/pathways. B) Schematic workflow of combination therapy (CT) resistant cells selection: A375N cells were maintained under continued combination therapy (CT) exposure (Dabrafenib 250 nM and Trametinib 10 nM); after a minimal residual disease (MRD) phase of quiescence, proliferating A375DR cells are expanded. Dot plot showing the gene sets significantly enriched in A375DR cells compared to A375N cells, detected by GSEA using the KEGG database, considering RNA-seq. Results are split into activated and suppressed processes/pathways. C) Immunoblot analysis of mTORC1 activation markers (p-S6K, p-S6), ribosomal proteins (S6, RSP3, RPL23, RPL26), and Erk1/2 activation (p-Erk1/2) inA375N and A375DR cells, showing mTORC1 blockade, ERK1/2 inhibition, and modulation of ribosome biogenesis D) Polysome profiles of A375N and A375DR cells showing the distribution of ribosomal subunits (40S and 60S), monosomes (80S), and actively translating polysomes. Quantification of the area under each peak was performed to assess differences in ribosome assembly and translational activity between the two cell lines. Data are presented as mean ± SD, and statistical significance was determined using an unpaired t-test.

To better investigate the role of CT in BRAFV600E melanoma, we also generated an acute model of resistance by treating A375N cells with the CT of Dabrafenib 250 nM and Trametinib 10 nM for 72 h (A375+CT). We next characterized how CT impacts on melanoma cell proliferation. A375DR cells and showed a reduced doubling time and slower proliferation rate than the naïve counterpart (Figure S1C), along with an altered cell-cycle composition (Figure S1D). In particular, both chronically and acutely treated cells (A375DR and A375N + CT) showed increased fraction of cells in the G0/G1 phase (78.3% and 77.6% respectively compared to 44.6% in A375N), and decreased fraction of cells in the G2M phase (11% and 13.8% compared to 26.6% in A375N) and S phase (10.3% and 8.6% compared to 28.2% in A375N), with no substantial modulation of cell fraction in the SubG1 phase.

To further assess the mechanism underlying the suppressed ribosome/translation machinery led by CT, we investigated the protein synthesis rate in our CT model using a puromycin pulse assay (Figure S1E,F). We confirmed that in both acute and chronic treatment, melanoma cells display a significant reduction in global protein synthesis and mRNA translation. Such a decrease is accompanied by weakened mTORC1 signaling in both acute and chronic settings (Figure 1C and S1G). Indeed, mTORC1 is directly targeted by the activated MAPK/ERK pathway. We were able to recapitulate the same effect of the CT in a mouse BRAFV600E melanoma model by using YUMM1.7 cells exposed to CT (Meeth *et al*, 2016)(Figure S1H,I).

To better characterize mRNA translational modulation in A375DR cells, we conducted polysome profiling, a technique that separates ribosomal subunits, monosomes, and polysomes/mRNA bound by multiple translating ribosomes by sedimentation velocity through a sucrose density gradient, allowing us to infer global translation activity from the relative distribution of mRNA across these fractions (Chassé *et al*, 2017; Gandin *et al*, 2014). This analysis revealed a marked remodeling of ribosome distribution in A375DR cells compared to A375N. Specifically, although the 40S and 60S ribosomal subunit fractions remained unchanged, both the abundance of polysomes and the proportion of ribosomes engaged in polysomes were significantly reduced (Figure 1D). Conversely, the free 80S ribosome fraction was increased, indicating an accumulation of non-translating ribosomes and an overall reduction in translational activity. Notably, this profile closely resembles that observed following mTORC1 pathway inhibition, which is characterized by polysome disassembly and increased 80S monosome accumulation (Shetty *et al*, 2023; Thoreen *et al*, 2012).

Overall, we found that combination of MAPKi and MEKi treatment in BRAF-mutated human melanoma and cells downregulates mTORC1 activation and global mRNA translation, decreasing protein synthesis and proliferative potential despite preserved cell viability. We might argue that targeted therapy acts to foster new adaptation for promoting selective protein synthesis that, at least partially, bypasses mTORC1 dependence and rewires its translation programs.

### CT-resistant cells translationally boost primary cilia formation

We next interrogated bulk RNA sequencing data of A375DR melanoma cells to identify novel compensatory mechanisms that may promote treatment resistance. We performed differential gene expression analysis (Figure 2A) followed by an over-representation analysis (ORA) (Figure 2B) of significantly upregulated genes. These analyses revealed an enrichment in gene sets related to “primary cilium” and “cilium assembly”, indicating an upregulation of cilia-associated processes in A375DR cells. Additionally, we discovered that some genes involved in cilia disassembly were significantly downregulated (*AURKA, CDK1, NEK2, NDE1, PLK1*)(Wang *et al*, 2013) (DeVaul *et al*, 2017; Korobeynikov *et al*, 2017; Liang *et al*, 2016). These findings suggest a mechanistic link between BRAFi and MEKi treatment and ciliogenesis in melanoma CT resistance.

**Figure 2.**
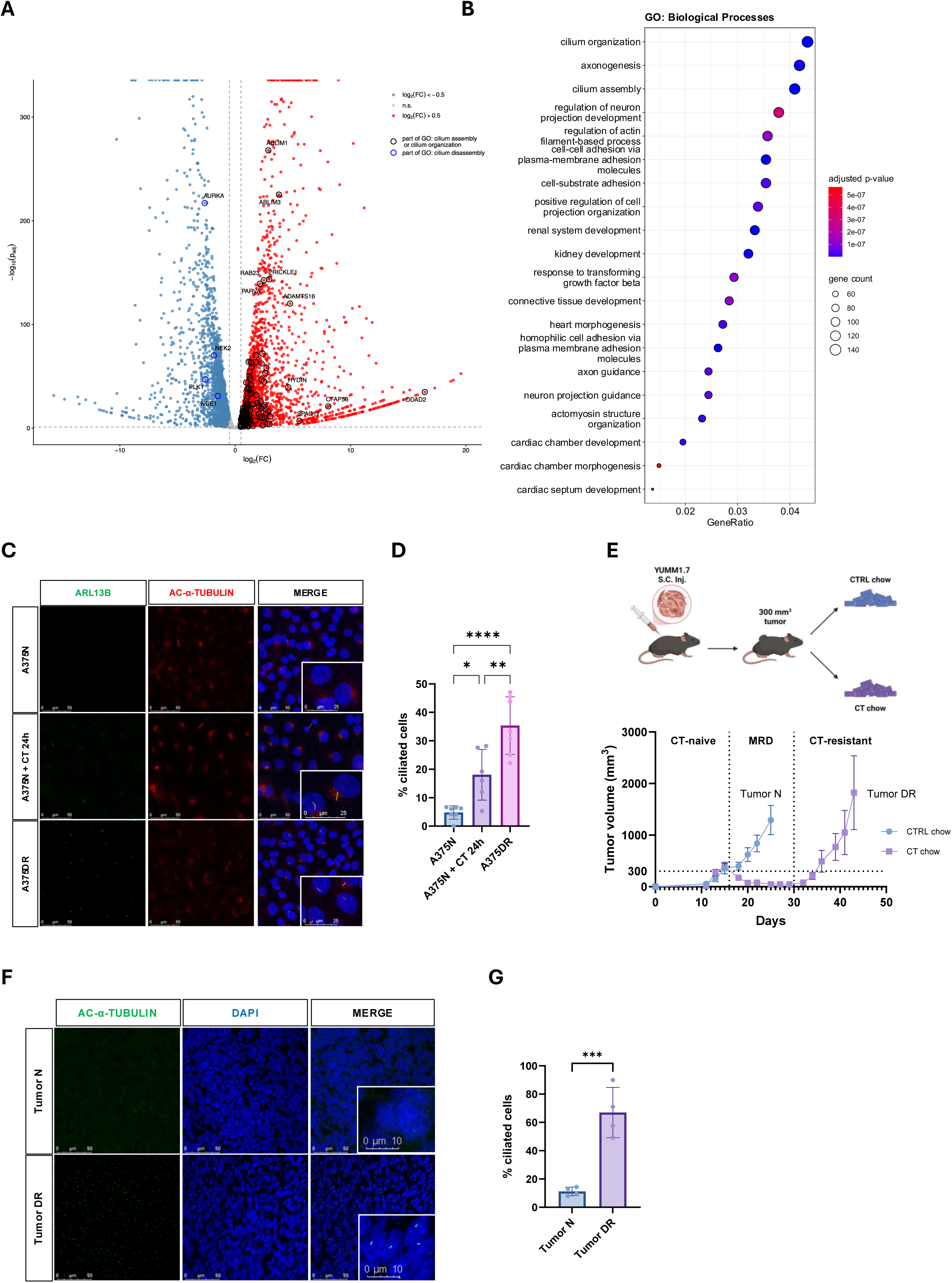
CT-resistant cells are characterized by primary cilia formation. A) Volcano plot showing differential gene expression between A375DR and A375N cells. Significantly changing genes belonging to the GO-BP terms “cilium assembly” and “cilium organisation” are marked with a black circle, while genes changing significantly belonging to the GO-BP term “cilium disassembly” are marked with a blue circle. Dotted lines indicate the threshold values for the log2(FC) and the adjusted p-value. B) Dot plot showing the gene sets significantly enriched in A375DR and A375N cells detected by over-representation analysis (ORA) using the Gene Ontology:Biological processes database, considering RNA-seq. C) Immunofluorescence staining of primary cilia in A375N cells, A375DR cells, and A375N cells treated for 24 h with combination therapy (CT; dabrafenib 250 nM + trametinib 10 nM), using antibodies against ARL13B and acetylated α-tubulin (ac-α-TUBULIN). Nuclei were counterstained with DAPI. D) Quantification of the percentage of ciliated cells from the experiment in (C); data are shown as mean ± SD; statistical significance was calculated by one-way ANOVA with Tukey’s multiple comparisons test E) Schematic workflow for the generation of CT-resistant tumors *in vivo*, and tumor growth of YUMM1.7 cells. YUMM1.7 cells were injected subcutaneously into the flank of C57BL/6N mice. When tumors reached ∼300 mm³, mice were fed with control chow or chow containing Dabrafenib and Trametinib continuously until the experimental endpoint. Tumor volume was measured three times a week. Data are shown as mean ± SEM (n = 4 mice per group). F) Immunofluorescence staining of primary cilia (ac-α-TUBULIN) in sections of YUMM1.7 tumors from the experiment in Figure 1B. Nuclei were counterstained with DAPI G) Quantification of the percentage of ciliated cells in the tumors shown in (F); data are shown as mean ± SD. Statistical significance was calculated by unpaired t-test.

To investigate the mechanism involved in ciliogenesis in the context of globally reduced protein synthesis, we used Ribo-seq data to compare translated mRNAs in A375DR with A375N cells (Clamer *et al*, 2018). We found that many genes related to ciliogenesis were increasingly translated in A375DR compared to A375N cells by examining ribosome occupancy in transcriptionally upregulated genes from the cilia-related GO-BP terms. These transcripts showed increased occupancy at both the CDS (*ATAT1*, *ODAD2, BBS4, DNAH5, PKD2, TMEM237*) but also at 5’UTR (*ODAD2, CFAP263, BBS4*) (Figure S2A,B).

Thus, to further investigate the mechanism underlying increased ciliogenesis, we also measured active translation of *ATAT1, TMEM237, BBS4, PKD2* mRNAs in polysomes vs. the subpolysome fraction from A375DR vs. A375N cells (Figure S2C). We discovered that they are enriched in the polysome fractions in A375DR compared to A375N, supporting their active translation in CT-resistant melanoma cells. We confirmed that other mRNAs showing no transcriptional modulation (e.g., *ITGA2, ITGA5*) did not show the same behavior in the polysomal fraction distribution, suggesting a cilia-specific translational program.

A375 is conventionally regarded as poorly ciliated (Zingg *et al*., 2018), so to validate our observation, we assessed ciliation in A375DR cells. A375N melanoma cells showed very few ciliated cells, whereas the percentage of ciliated cells increased significantly after acute CT and, particularly, during resistance acquisition (Figure 2C,D). Notably, we observed no differences in cilia length across the experimental conditions (data not shown). We also exclude that cilia formation is dependent on drug-resistant influence on proliferation and cell cycle progression (Snedecor *et al*., 2015).

To test whether such therapy resistance-dependent increase in cilia occurs also in vivo, we developed a model of in vivo drug resistance using the C57BL/6J syngeneic mouse melanoma cell line YUMM1.7 (Meeth *et al*., 2016). Cells were subcutaneously injected, and mice received CT-containing chow once the tumor reached 300 mm³ in volume. This approach mimicked clinical target therapy resistance, showing an initial rapid tumor shrinkage, followed by a plateau phase of minimal residual disease (MRD), and ultimately, tumor regrowth despite ongoing drug treatment (Figure 2E). We next isolated tumors from mice under normal chow (CTRL chow: tumor N) and tumors from mice under CT chow (CT chow: tumor DR). Tumor DR again showed a markedly increased frequency of ciliated cells compared to tumor from control conditions (Figure 2F,G).

We conclude that adaptation to CT treatment leads to ciliogenesis in melanoma BRAF-mutated cells and tumors. Because primary cilia depend on various structural proteins and enzymes (Reiter & Leroux, 2017), we showed that ciliogenesis in melanoma adaptation to CT likely involves coordinated regulation across transcriptional and translational responses, possibly coordinated by unforeseen therapy resistance and adaptation mechanisms.

### Primary cilia are functionally relevant in targeted therapy resistance development

To assess the role of primary cilia in CT therapy resistance, we generated a genetic model of cilia abrogation by knocking out Intraflagellar Transport 88 (*IFT88*) gene, which has been shown to be indispensable for primary cilia formation (Haycraft *et al*, 2007). Using CRISPR/Cas9 system, we generated single A375N clones and selected the two showing complete gene knockout (clones #1 and #6)(Figure S3A). We then tested these clones for cilia abrogation by acute exposure to CT. While control (gSCR) cells responded to MAPKi+MEKi with a significant increase in ciliation, we detected no cilia in the clones exposed to the same stimulus, confirming cilia abrogation (Figure 3A,B). Importantly, we detected no reduction in proliferation rate and no differences in global cell viability following *IFT88* knockout in both clones (Figure S3B,C). To test whether the presence of cilia might influence sensitivity to MAPK and MEK inhibitors, we treated *IFT88*^KO^ melanoma cells with CT. *IFT88*^KO^ melanoma cells showed increased sensitivity to CT, with a strong reduction in cell proliferation in both acute and chronic CT (Figure 3C, D), along with a significant decrease in IC50 (Figure 3E) and clonogenicity (Figure 3F).

**Figure 3.**
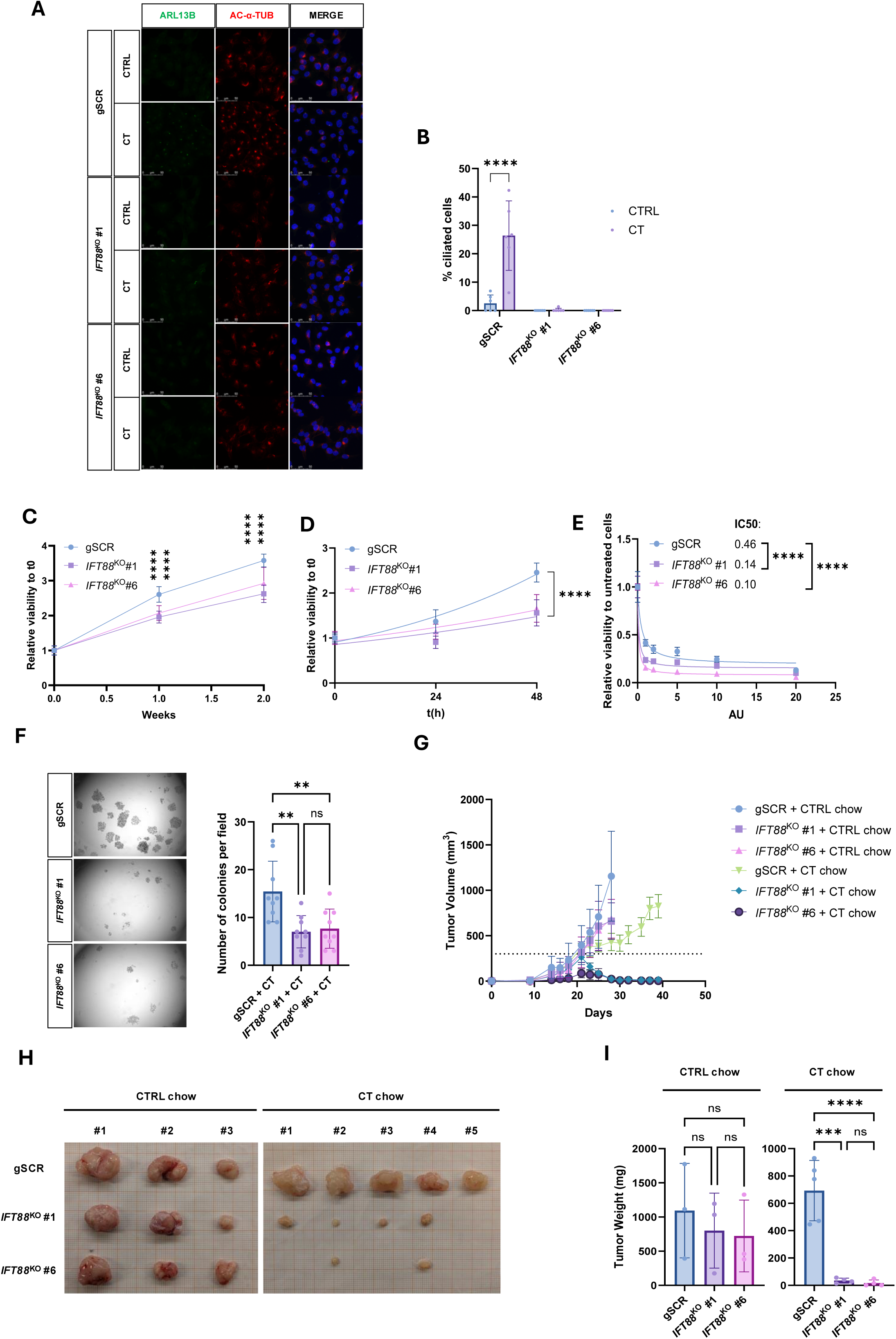
Primary cilia are functionally relevant in CT resistance adaptation. A) Immunofluorescence staining of primary cilia with ARL13B (green) and acetylated α-tubulin (ac-α-TUBULIN, red) in A375N scrambled control (sSCR) and *IFT88* knockout (KO) clones treated for 24 h with combination therapy (CT; dabrafenib 250 nM + trametinib 10 nM). Nuclei were counterstained with DAPI. B) Quantification of the percentage of ciliated cells from the experiment in (A); statistical significance was calculated by two-way ANOVA with Tukey’s multiple comparisons test. C) Proliferation assay of A375N gSCR and IFT88^KO^ clones under CT over two weeks. Cells were stained with crystal violet at the indicated time points and quantified with ImageJ; proliferation is expressed as relative viability compared to time 0h; statistical significance was calculated by one-way ANOVA with Tukey’s multiple comparisons test at each time point. D) Proliferation assay of A375N gSCR and IFT88^KO^ clones under CT for 72 h, stained with crystal violet and quantified with ImageJ. Curves were fitted with an exponential (Malthusian) growth model by least squares regression, and doubling times were compared between curves by extra sum-of-squares F test. E) Dose-response assay of A375N gSCR and IFT88^KO^ clones treated for 72 h with increasing concentrations of CT (1 AU = dabrafenib 50 nM + trametinib 2 nM). Cell viability was assessed by crystal violet staining, quantified with ImageJ and expressed relative to untreated cells. Dose-response curves were fitted with the “[Inhibitor] vs. response (three parameters)” model by least squares regression, and IC50 values (in AU) were compared between clones by extra sum-of-squares F test. F) Colony formation assay of A375N gSCR and IFT88^KO^ clones cultured for 14 days under CT. Colonies were stained with crystal violet; representative wells are shown together with the quantification obtained with ImageJ (bar plot). Statistical significance was calculated by one-way ANOVA with Tukey’s multiple comparisons test. G) Tumor growth of A375N gSCR and IFT88^KO^ A375 clones injected subcutaneously into NSG mice (5 × 10ff6 cells per injection). When tumors reached ∼300 mm³ in size, mice were fed control chow or chow containing dabrafenib and trametinib. Tumor volume was measured three times a week. Data are shown as mean ± SEM. H) Representative pictures of the tumors excised at endpoint from the experiment in (G). I) Tumor weight at endpoint from the experiment in (G). Statistical significance was calculated by one-way ANOVA with Tukey’s multiple comparisons test.

To test whether this increased sensitivity of cilia-depleted melanoma cells to CT also occurs in vivo, we subcutaneously injected gSCR cells and *IFT88*^KO^ A375 clones into immunocompromised NOD-SCID-gamma mice (NSG) and started treating the mice with chow containing CT once the tumor reached a palpable size of 300-500 mm^3^. Untreated gSCR and *IFT88*^KO^ tumors showed no differences in growth rate or weight (Figure 3 G,H, I), highlighting the dispensability of cilia under A375N conditions. In contrast, when we exposed the injected mice to CT, the tumors responded strikingly differently. gSCR cells showed slowed growth and tumor weight, as expected, while KO cells shrank rapidly after CT treatment, and nearly no tumor was visible at the endpoint. Notably, two mice injected with one clone experienced complete tumor regression (Figure 3 H,I).

We conclude that genetic ablation of cilia does not affect “naïve” untreated BRAFV600E melanoma cells, in which cilia appear to be non-essential. However, the lack of ciliogenesis markedly increases sensitivity of BRAFV600E melanoma cells and tumor to CT both in vitro and in vivo, suggesting that cilia are essential during acquired CTresistance in melanoma. Therefore, primary cilia are probably not merely an accessory of drug-resistant cells but are actively implicated in resistance acquisition. These findings raise the possibility of a mechanistic link between CT treatment and ciliogenesis in melanoma, in which cilia may serve as an adaptive mechanism in response to MAPK pathway inhibition.

### CT-resistant melanoma cells reprogram glutamine and aspartate metabolism

Primary cilia have been linked to cellular and mitochondrial metabolism (Miller *et al*, 2025; Oh *et al*., 2015), particularly adipogenesis {Hilgendorf, 2024 #34} and glutaminolysis (Steidl *et al*., 2023). Both pathways, along with global metabolic rewiring, have been functionally implicated in the target therapy-resistant phenotype in melanoma (Aloia *et al*, 2019; Haq *et al*., 2013; Vergani *et al*, 2022). We therefore question whether primary ciliogenesis in A375DR cells may help sustain such metabolic rewiring. To investigate this, we first characterized the metabolic state of A375DR cells to eventually assess its functional connection to primary cilia. At first, we measured basal and maximal respiration of A375DR and found that it was substantially increased compared to A375N (Figure S4A). We also observed decreased Extracellular Acidification Rate (ECAR) consistent with reduced glycolysis in A375DR cells (Baenke *et al*, 2016; Parmenter *et al*, 2014). To provide a comprehensive overview of metabolic reprogramming in A375DR cells, we conducted a targeted metabolomic profiling comparing A375N and A375DR cells. The analysis revealed significantly distinct metabolic profiles, highlighting the extensive metabolic reprogramming that underpins the phenotypic transition during CT resistance (Figure 4A). Among the most increased amino acids, we found glutamine and aspartate levels, but not glutamate (Figure 4B). While increased aspartate levels have never been reported, glutamine dependence and increased glutamine uptake have been shown in melanoma cells treated with MAPK pathway inhibitors, possibly promoting glutaminolysis and mitochondrial oxidative metabolism (Hernandez-Davies *et al*, 2015). Accordingly, such metabolic rewiring is accompanied by reduced glycolysis (Figure S4A)(Baenke *et al*., 2016; Wang *et al*, 2018). Our results also showed that A375DR cells displayed reduced overall ATP charge and reduced ribo- and deoxyribonucleotide and pyrimidine levels, which may indicate decreased global energy demand and nucleotide synthesis due to slower cellular proliferation (Figure S4B). Generally, metabolites involved in the TCA cycle were not significantly altered (Figure S4C).

**Figure 4.**
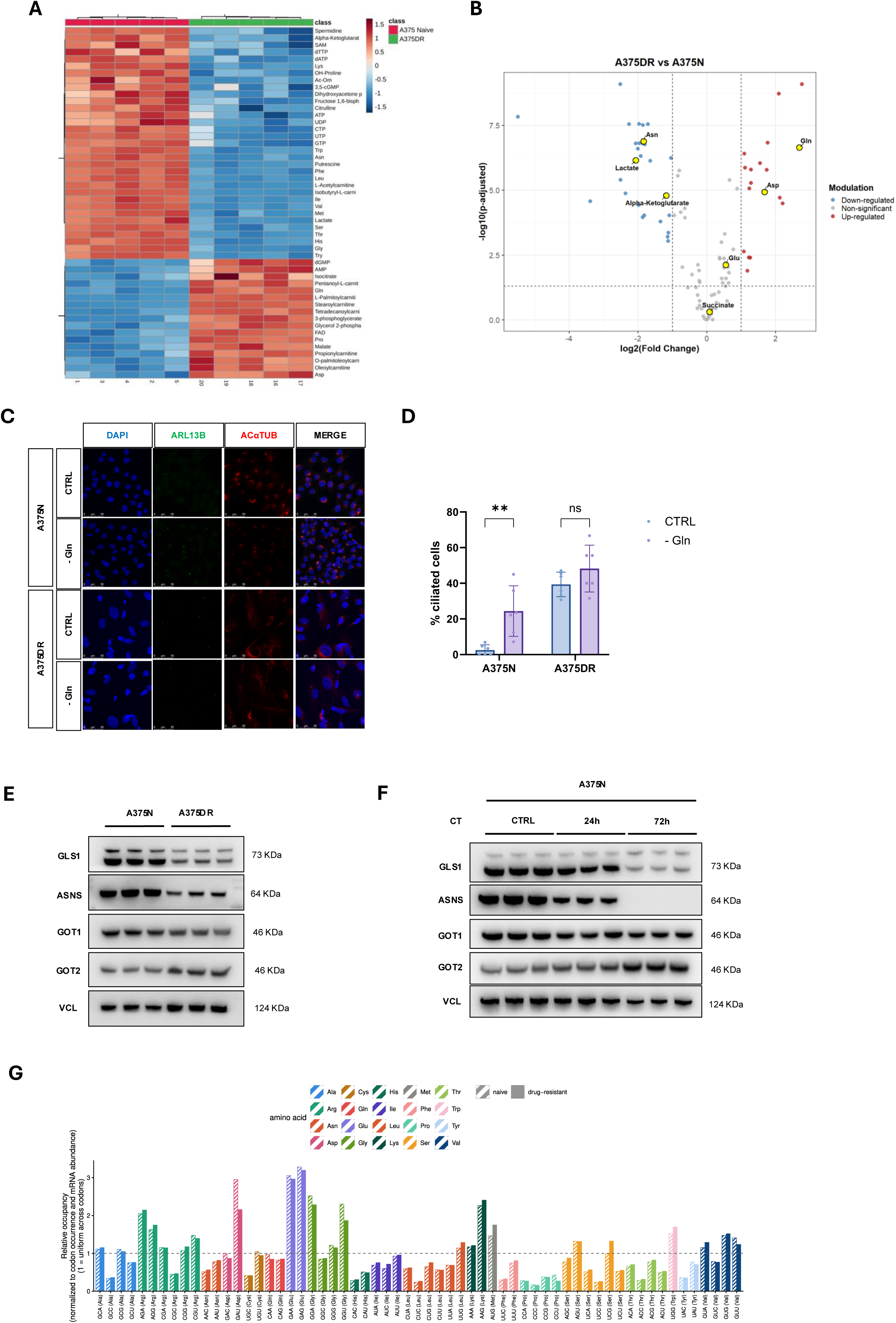
CT-resistant cells display Glutamine and Asp metabolism reprogramming. A) Heatmap of the top 50 metabolites differentially abundant between naïve and A375DR cells. Metabolomic data were normalized by median, log-transformed and Pareto-scaled in MetaboAnalyst. Hierarchical clustering of metabolites (rows) and samples (columns) was performed using Euclidean distance and Ward’s linkage. The color scale represents scaled metabolite abundance, from low (blue) to high (red). B) Volcano plot of metabolites differentially abundant between A375DR and A375N cells, showing log2(fold change) (DR vs naïve) versus −log10(adjusted p-value). Fold changes were calculated as ratios between group means (DR/naïve) on median-normalized data prior to log transformation and scaling. P-values were calculated by unpaired t-test on log-transformed data and adjusted for multiple testing by the Benjamini–Hochberg FDR method. Metabolites with |log2 fold change| > 1 and p adj < 0.05 were considered significantly upregulated or downregulated. C) Immunofluorescence staining of primary cilia with ARL13B (green) and acetylated α-tubulin (ac-α-tubulin, red) in naïve and A375DR cells after 24 h of glutamine (Gln) starvation (control condition: complete medium with 4 mM glutamine). Nuclei were counterstained with DAPI. D) Quantification of the percentage of ciliated cells from the experiment in (C). Data are shown as mean ± SD. Statistical significance was calculated by two-way ANOVA with Tukey’s multiple comparisons test. E) Immunoblot analysis of glutamine-aspartate metabolic enzymes (GLS1, GOT1,and GOT2), and asparagine synthetizing enzyme (ASNS), in lysates of A375N and A375DR, showing reduced expression of GLS1, ASNS and ATF4, and increased expression of aspartate synthesizing enzyme GOT1. F) Immunoblot analysis of glutamine-aspartate metabolic enzymes (GLS1, GOT1, GOT2), and asparagine synthetizing enzyme (ASNS) in A375N cells treated for 24 or 72 h with combination therapy (CT; dabrafenib 250 nM + trametinib 10 nM). G) Codon-resolved ribosome occupancy in A375N and A375DR cells. Bars show relative ribosome occupancy (normalized to the codon frequency and RNA abundance) at each of the 61 sense codons, grouped by amino acid.

It has been reported that resistant cells, being less dependent on glucose for proliferation, become more addicted to glutamine, the second most important alternative carbon source for tumor growth (Ding *et al*, 2026). To investigate the role of increased glutamine in A375DR cells, we treated these cells with the GLS1 inhibitor CB839 and evaluated cell viability (Figure S4D). A375DR cells showed higher sensitivity compared to naive cells to glutaminolysis inhibition. We also treated cells with the glutamine antagonist, 6-Diazo-5-oxo-L-norleucine (DON), which inhibits other glutamine-using enzymes, amino acid transporters, and transglutaminases (Figure S4E). Here we discovered that A375N cells are more sensitive to this glutamine antagonist, suggesting a completely different way of using glutamine between A375DR vs A375N cells. Next, we examined the impact of glutamine deprivation in our melanoma cell model, noting a significant increase in the proportion of ciliated cells among naïve cells, whereas the DR counterpart, which already exhibits high levels, showed no significant change (Figure 4C,D). This observation supports a functional relationship between glutamine availability and primary cilia formation in our melanoma model. Notably, although previous studies have attributed this connection to ASNS localization at the ciliary basal body (Steidl *et al*., 2023), our findings reveal a substantial reduction in ASNS expression, implying an alternative mechanism.

Interestingly, we discovered a strong accumulation of aspartate in CT-resistant BRAFV600E melanoma cells. This was quite unexpected and led us to investigate possible reasons. At first, we explored expression levels of critical metabolic enzymes related to aspartate synthesis and usage. We noticed that in both acute CT treatment and resistance acquisition, A375DR melanoma cells are characterized by increased aspartate-synthesizing enzyme GOT2 expression but not GOT1 (Figure 4E,F). Modulation of GOT2 expression appears to be part of a broader rewiring of glutamine and aspartate metabolism, as major other metabolic enzymes involved display consistent downmodulation, especially GLS1 and ASNS. Next, we assessed whether the elevated aspartate levels might influence translational processes in A375DR cells. Thus, we measured codon-resolved ribosome occupancy. Ribo-seq data allow codon-resolved ribosome occupancy to be measured, which serves as a proxy for the time ribosomes spend decoding each codon (Dana & Tuller, 2014). We therefore calculated ribosome occupancy at each of the 61 sense codons, normalized to the frequency of the codon in the coding sequences and to the transcript abundance. We then normalised relative to the average codon, such that a value of 1 indicates uniform occupancy across codons (Figure 4G). Ribosome occupancy at the aspartate codon GAU was lower in A375DR than in A375N cells, implying faster decoding of this codon in A375DR cells. Interestingly, most aminoacyl-tRNA synthetases, including *DARS1* and *DARS2*, show negative transcriptional modulation in A375DR cells (Figure S4F), suggesting that the decrease in aspartate codon occupancy does not depend on these enzymes’ activity but rather on increased aspartate availability.

Overall, our A375DR model recapitulated some of the metabolic features of therapy resistance reported in previous studies, including decreased glycolytic flux and increased OXPHOS (Corazao-Rozas *et al*., 2013). Our data suggest that A375 cells with acquired CT resistance are more reliant on glutamine dependency (Baenke *et al*., 2016). At the same time, our data on A375DR converge on increased intracellular aspartate levels, which do not appear to affect anabolic or catabolic mechanisms but seem to influence translation via an unknown mechanism.

### GOT1/GOT2 enzymes and aspartate levels are not related to CT-dependent primary ciliogenesis

Aspartate metabolism is a vital cellular process in which aspartate acts as a central hub for building proteins, creating DNA/RNA building blocks, moving energy inside the cell and activating mTOR signaling. Most cell types and cancer types cannot import aspartate from the extracellular environment; therefore, they must produce it intracellularly. Modulation of glutamate-oxaloacetate transaminase 1 (in the cytosol) (*GOT1*) and 2 (in the mitochondria) (*GOT2*) expression has been shown to directly affect intracellular aspartate levels, including in the context of cancer (Rossi *et al*, 2022; Yang *et al*, 2018). To define the role of aspartate accumulation in A375DR adaptation to CT, we knocked down (KD) *GOT1* and *GOT2* in A375N and A375DR cells and studied modulation of translation, ciliogenesis, and drug sensitivity (Figure S5A). Genetically silencing *GOT1* and *GOT2* has been used as a direct approach to control aspartate levels with *GOT1* silencing raises aspartate levels by reducing its consumption, whereas *GOT2* silencing lowers asp levels by hindering its synthesis (Brunner *et al*, 2026).

At first, we evaluated global variation in global protein synthesis in A375N and A375DR cells under acute CT treatment (Figure 5A, S5B). Interestingly, a puromycin pulse showed that, upon *GOT1* or *GOT2* KD, A375Ncells had an increased translation rate, which may be partially explained by increased aspartate availability. However, we didn’t observe the same effect in cells under acute or chronic CT treatment. Also, neither KD affected cell proliferation in both naïve and DR conditions (Figure 5B-C). In addition, we evaluated drug-sensitivity variation upon *GOT1* or *GOT2* KD, but we didn’t observe significant variation in naïve cells or restored sensitivity in DR cells (Figure 5D-E).

**Figure 5.**
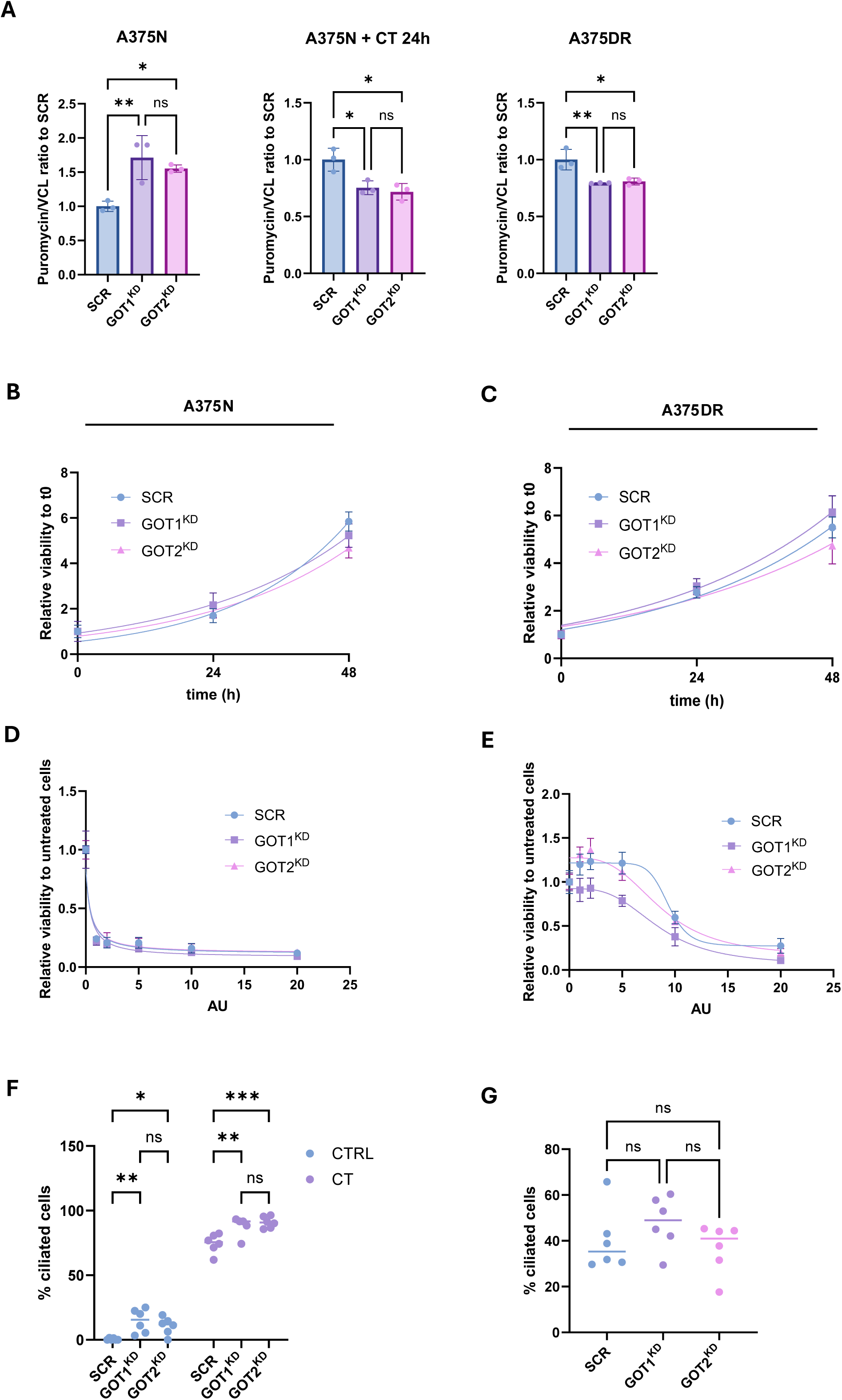
Aspartate and GOT do not àect primary cilia synthesis and drug sensitivity. A) Quantification of global protein synthesis by puromycin pulse assay in A375N cells, A375N cells treated for 24 h with combination therapy (CT; dabrafenib 250 nM + trametinib 10 nM), and A375DRcells, upon GOT1 or GOT2 knockdown (KD) or scrambled (SCR) control. Data are shown as mean ± SD. Statistical significance was calculated by one-way ANOVA with Tukey’s multiple comparisons test. B) Growth curves of A375N cells with GOT1 or GOT2 KD (and SCR), stained with crystal violet at time 0, 24 and 48 h and quantified with ImageJ. Proliferation is expressed as relative viability compared to t0. Curves were fitted with an exponential (Malthusian) growth model by least squares regression using GraphPad Prism, and doubling times were compared between curves by extra sum-of-squares F test. C) Growth curves of A375DR cells with GOT1 or GOT2 KD (and SCR), as described in (B). D) Dose-response assay of A375N cells with GOT1 or GOT2 KD (and SCR) treated for 72 h with increasing concentrations of CT (1 AU = dabrafenib 50 nM + trametinib 2 nM). Cells were stained with crystal violet, quantified with ImageJ and expressed relative to untreated cells. Dose-response curves were fitted with the “[Inhibitor] vs. response (three parameters)” model by least squares regression, and IC50 values were compared between conditions by extra sum-of-squares F test. E) Dose-response assay of A375DR cells with GOT1 or GOT2 KD (and SCR), as described in (D). F) Quantification of the percentage of ciliated cells from immunofluorescence images of A375N cells with GOT1 or GOT2 KD (and SCR), untreated or treated for 24 h with CT. Statistical significance was calculated by two-way ANOVA with Tukey’s multiple comparisons test G) Quantification of the percentage of ciliated cells from immunofluorescence images of A375DR cells with GOT1 or GOT2 KD (and SCR). Statistical significance was calculated by one-way ANOVA with Tukey’s multiple comparisons test.

Subsequently, we examined changes in ciliogenesis after *GOT1* or *GOT2* KD in A375N cells subjected to acute CT treatment (Figure 5F). Both *GOT1* and *GOT2* KDs significantly increased the proportion of ciliated cells in naïve populations under basal conditions and after acute CT exposure. The finding that both knockdowns increase ciliogenesis in A375N independently of CT treatment suggests that this effect is independent of CT-dependent aspartate increase and may reflect a broader metabolic stress response. Accordingly, when we examined changes in ciliogenesis after *GOT1* or *GOT2* KD in A375DR cells, we did not score significant differences (Figure 5G).

Overall, although GOT2 levels and aspartate accumulation characterized CT-adaptation in melanoma, they do not appear to directly affect primary ciliogenesis or impact global protein translation, leaving its functional role undetermined.

### Metabolic insights into the role of primary cilia in drug resistance

After establishing that increased primary cilia play a functional role in therapy resistance and observing their functional connection to glutamine metabolism, we next explore how primary cilia contribute to the development of resistance possibly via metabolic adaptation.

At first, we conducted targeted metabolomic profiling of A375N IFT88^KO^ cells (Figure S6A). *IFT88*^KO^ does not appear to affect global cellular metabolism in BRAF-mutated melanoma cells. In detail, *IFT88*^KO^ clones exhibited a significantly increased succinate/α-KG ratio and a decrease in fatty acid β-oxidation, although the levels of glutamine, glutamate, and α-KG did not change significantly (Figure S6B).

We then investigated the metabolic function of primary cilia under A375N conditions by depriving cells of glutamine, a process we have shown enhances ciliogenesis (Figure 4C,D). We observed no substantial differences in cell viability when comparing cilia-deprived cells after 24 hours of Glutamine starvation compared to controls (Figure 6A,B), suggesting that the IFT88^KO^ cells may compensate for cilia loss by activating alternative pathways. Protein expression analysis showed that IFT88^KO^ cells express glutamine synthetase (GS) under normal conditions, whereas GS remains undetectable in control (gSCR) cells (Figure 6C). Most notably, GS expression is markedly enhanced during glutamine starvation in IFT88^KO^ cells, suggesting a potential mechanism that may bypass the cilia-mediated response to glutamine deprivation.

**Figure 6.**
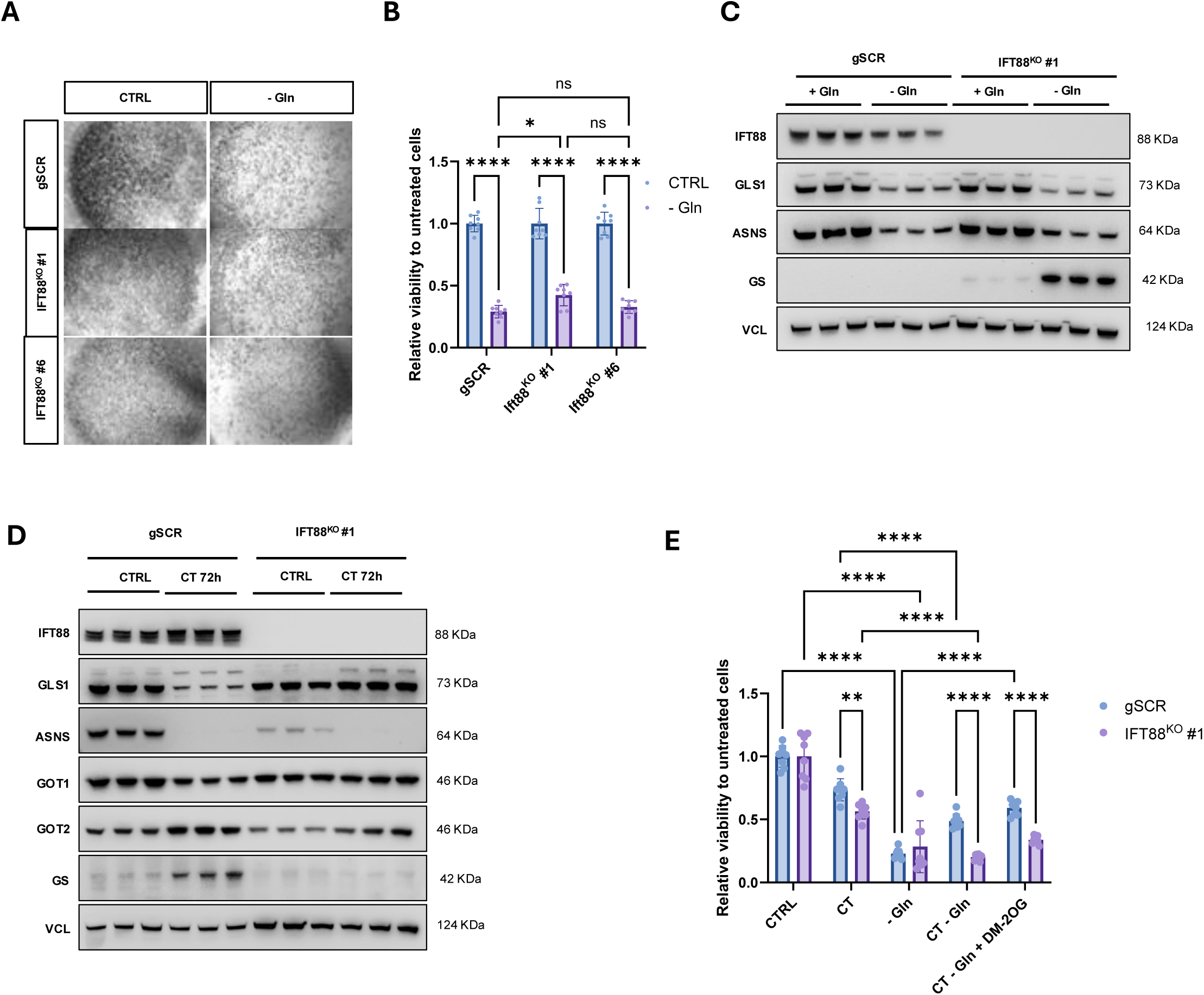
Metabolic insight into the role of primary cilia in CT resistance. A) Representative images of A375N scrambled control (SCR) and *IFT88* knockout (KO) A375 clones after 24 h of glutamine (Gln) starvation or in control medium (4 mM glutamine). Cells were stained with crystal violet. B) Quantification of the experiment in (A), expressed as relative viability compared to the control medium condition of each cell type. Cell viability was assessed by crystal violet staining, quantified with ImageJ. Data are shown as mean ± SD. Statistical significance was calculated by two-way ANOVA with Šídák’s multiple comparisons test. C) Immunoblot analysis in gSCR and IFT88^KO^ #1 A375 cells of IFT88, GLS1, ASNS, and GS expression upon glutamine starvation. D) Immunoblot analysis of IFT88, GLS1, GOT1, GOT2, ASNS, and GS expression in A375N gSCR and IFT88^KO^ A375 cells treated for 72 h with combination therapy (CT; dabrafenib 250 nM + trametinib 10 nM) or vehicle. E) Viability assay of A375N scrambled control (gSCR) and *IFT88* knockout (KO) A375 clones treated for 72 h with combination therapy (CT; dabrafenib 250 nM + trametinib 10 nM), glutamine (Gln) starvation, cell-permeable dimethyl oxoglutarate (DM-2OG; 2 mM), or the indicated combinations. Cells were stained with crystal violet and quantified with ImageJ. Viability is expressed relative to the control condition (complete medium, vehicle) of each cell type. Data are shown as mean ± SD. Statistical significance was calculated by two-way ANOVA with Šídák’s multiple comparisons test.

Because we could not generate drug-resistant *IFT88*^KO^ melanoma cells, we treated *IFT88*^KO^ melanoma cells with CT and analyzed glutamine and aspartate enzymes (Figure 6D). Acute treatment recapitulated A375DR behavior with a decrease in GLS1 and ASNS expression and an increase in GOT2 and GS expression (Figure 4F). Interestingly, some of these metabolic enzyme switches were lost in *IFT88*^KO^ cells, such as GLS1 decrease (*IFT88*^KO^ cells keep expressing GLS1 after therapy treatment) and GS is no longer induced by therapy treatment if cilia are not present. ASNS remains unchanged in all conditions, suggesting that cilia are not regulating this enzyme in melanoma (Figure 6D).

To better define the metabolic mechanism involved in cilia-mediated control of metabolic rewiring in target therapy conditions, we treated *IFT88*^KO^ A375N cells with CT in combination with glutamine deprivation (Figure 6E). *IFT88*^KO^ cells are more sensitive to CT alone, but do not differ during glutamine starvation, possibly because *IFT88*^KO^ cells induce GS (Figure 6C). Under Glutamine starvation and in the presence of BRAFi and MAPKi, A375N gSCR cells are more viable compared to IFT88^KO^ cells, suggesting a putative function for cilia in regulating glutamine uptake that in turn supports metabolic adaptation under CT (Figure 6E). Interestingly, CT can promote GS induction in control cells, which may further increase glutamine levels, but this does not happen in cilia-depleted A375N cells. When cells are treated with Dimethyl 2-oxoglutarate (DM-2OG) to fuel the TCA cycle, neither cell line appears affected, suggesting that cilia-dependent glutamine dependency may have an additional, unforeseen function in A375 cells.

Since the role of primary cilia in melanoma drug resistance is not fully understood, it is crucial to consider other potential functions, given cilia’s diverse regulatory roles. Among the signaling pathways examined in the introduction, the SHH pathway is of particular interest because of its tight connection to primary cilia formation and dismantling, as well as its role in other cancer types (Bangs & Anderson, 2017; Han *et al*, 2009; Wang *et al*, 2010). Indeed, we assessed expression of the SHH pathway effectors GLI1 and GLI2 using our bulk RNA sequencing data. The activation of the SHH pathway is supported by results of ORA of transcriptionally upregulated genes (significant enrichment of “Hedgehog signaling pathway” (KEGG) and “smoothened signaling pathway” (GO-BP), (see supplement data). Notably, *GLI2* showed a significant increase at the transcriptional level, whereas *GLI1* remained unchanged (Figure S6C). Consequently, we examined sensitivity to pathway inhibition in both naïve and DR cells. Interestingly, A375DR cells were significantly more sensitive to GANT-61, a selective inhibitor of GLI1 and 2 (Figure S6D). Overall, SHH pathway activity may serve as another way primary cilia contribute to resistance development. Instead of replacing metabolic activity, it likely acts as a parallel mechanism, since cilia perform numerous cell-specific functions based on their distinct composition (Collinson & Tanos, 2025; Wachten & Christensen, 2025).

## DISCUSSION

Despite recent progress, melanoma remains the most aggressive skin cancer, with therapy still limited due to acquired resistance. Melanoma’s aggressiveness and relapse are associated with tumor cells’ ability to dynamically adapt to new extracellular cues, including drug-dependent therapies. Melanoma cells can transiently and reversibly switch their phenotype, suggesting that, beyond genetic factors, transcriptional and translational mechanisms likely drive melanoma progression and therapy response. We discovered that one such mechanism is ciliogenesis. Genes involved in cilia assembly are importantly induced transcriptionally in A375DR cells. We also found that, beyond an overall switch in the mTOR/ribosome/translational machinery, these cilia genes are actively translated, suggesting that specific mRNA translational rewiring may be occurring in CT-treated melanoma cells. Further studies are needed to be able to identify whether specific eIFs or tRNA^aa^-specific mechanisms are involved (El-Hachem *et al*, 2024).

In cancer, the relationship between primary cilia and metabolism can vary considerably by context. Many cancers have fewer primary cilia or may lose them entirely (Chen *et al*, 2020; Kobayashi *et al*, 2017; Menzl *et al*, 2014). This reduction may reflect the inverse relationship between ciliogenesis and cell-cycle progression: primary cilia typically assemble in quiescent cells and disassemble before cells re-enter the cell cycle (Kim & Tsiokas, 2011; Plotnikova *et al*, 2008). However, ciliary loss in cancer is not always permanent. In response to treatment or genotoxic stress, some cancer cells can reassemble or lengthen their primary cilia, suggesting that ciliary regrowth may help them adapt to stress (Chen *et al*, 2021). In this setting, primary cilia may provide a specialized signaling platform that helps cancer cells sense extracellular stress and adjust survival or invasion pathways. In our experimental melanoma settings, we discovered a direct correlation between cilia function and metabolism adaptation. The capacity of primary cilia to influence metabolic adaptation is supported by the selective enrichment of receptors, ion channels, transporters, and signaling molecules within the ciliary compartment (Hansen *et al*, 2025; Wachten & Christensen, 2025). Primary cilia have been associated with resistance to a range of anticancer therapies. Resistant lung cancer cells often have more and longer primary cilia, which mediate several mechanisms of resistance to kinase inhibitors (Jenks *et al*., 2018). In pancreatic ductal adenocarcinoma, cisplatin-resistant cells regrow primary cilia through DNA damage-associated stress signaling; disrupting ciliogenesis makes these cells more sensitive to cisplatin. In triple-negative breast cancer, ciliary signaling associated with epithelial-to-mesenchymal transition (EMT) has also been linked to stem-like states and therapeutic resistance (Tessier *et al*, 2025). Together, these findings suggest that therapy- or stress-induced ciliogenesis can contribute to coordinate metabolic stress sensing, but also coordinate EMT-associated plasticity and pro-metastatic signaling (Jiang *et al*, 2014; Schlotfeldt *et al*, 2026; Wilson *et al*, 2021).

Our metabolic data comparing A375DR and A375N cells indicate that aspartate, proline and glutamine are the single largest amino acid changes in the set. Previous work has shown that increased glutamine uptake by CT leads to glutaminolysis, as CT-treated cells switch to OXPHOS instead of glycolysis (Baenke *et al*., 2016; Parmenter *et al*., 2014). Interestingly, in our model, glutamate increases only moderately, not significantly, while alpha-ketoglutarate is downregulated. Across these two steps, the pool shifts approximately 4.4-fold toward glutamine relative to glutamate and approximately 3.3-fold toward glutamate relative to alpha-ketoglutarate. These data suggest that DR cells have a break at the Glu-to-aKG step, with a partial block upstream as well. An interpretation is that A375DR cells import and retain glutamine but have largely stopped running it through canonical glutaminolysis into anaplerosis. Three non-exclusive mechanisms support this: reduced GLS1 flux with glutamine pooling up; GLUL induction converting glutamate and NH3 back to glutamine; or a shift of glutamine use away from carbon toward amide-nitrogen donation by the aminotransferases. Whether cilia are needed is important, and glutamine uptake is currently under investigation; we are also performing metabolic flux analysis to clarify glutamine use by A375DR cells.

An unforeseen result was the accumulation of aspartate in the CT-resistant cells. Cells make aspartate through transamination with glutamate-oxaloacetate transaminase 1 (in the cytosol) and 2 (in the mitochondria), which catalyze this reaction by transferring an amino group from glutamate to oxaloacetate. Alternatively, many cells might rely on mitochondrial glutamine catabolism to supply the carbon needed for this process (Ling *et al*, 2023). Aspartate may carry energy-carrying electrons across mitochondrial membranes to support cellular respiration in the malate-aspartate shuttle (aspartate availability influences engagement of the malate-aspartate shuttle). Aspartate is also directly involved in nucleotide synthesis: it donates atoms to build purines and pyrimidines, which form RNA and DNA. High aspartate levels activate the mTORC1 pathway, signaling the cell to grow and synthesize new proteins, which seems not to be the case here (Rae *et al*, 2025). However, aspartate cannot easily cross the outer cell membrane; as such, aspartate is an endogenous metabolic limitation for tumor growth (Sullivan *et al*, 2018). This poor permeability can limit cell growth when internal production slows. Under stress, such as low oxygen, some cells use specific surface transporters, like SLC1A3, to import aspartate from their environment (Sun *et al*, 2019). In our dataset, aspartate levels significantly increase in conjunction with a decrease in asparagine, resulting in roughly an 11.5-fold shift across the ASNS reaction. Aspartate accumulation is often interpreted as an indicator of electron transport chain competence, since aspartate synthesis relies on ETC-supported oxidation. Aspartate is typically a limiting factor for cell proliferation. In this context, it accumulates despite all nucleotide triphosphates declining and monophosphates increasing. The ATP/AMP ratio alone decreases by approximately 9.5-fold. This reflects a low-energy charge, reduced biosynthesis, and a slow-cycling state that represent the metabolic signature of a drug-tolerant *persister* cell rather than a rapidly proliferating resistant clone. Low asparagine matters in its own right: asparagine acts as an intracellular exchange factor for essential amino acid import. We showed that most of the essential amino acids are down (leucine, valine, phenylalanine, threonine, tryptophan, tyrosine, histidine, methionine, lysine). So low asparagine might explain global EAA depletion by impairment of EAA antiport. Accordingly, there is a similar corollary on the glutamine side: SLC7A5/LAT1 imports leucine by exporting glutamine. Glutamine trapped high inside with leucine depleted would suggest a stalled antiport. Low leucine also impairs mTORC1 signaling, which explains reduced global translation and low ATP demand. These findings point to an intersection between cilia and SLC (Solute Carrier) transporters, a crucial frontier in cellular signaling, homeostasis, and sensory physiology.

Overall, we investigate melanoma resistance to targeted therapy to identify new resistance mechanisms. We combined *in vivo* and *in vitro* approaches to demonstrate that cilia represent an unforeseen aspect of such resistance. Whether loss of primary cilia in melanoma resistance is causative, concomitant, or consequential to malignant transformation or invasion is still under discussion. Because of the pivotal role of primary cilia in hedgehog signaling (Bangs & Anderson, 2017), it will be interesting to determine whether the therapeutic potential of inhibiting the hedgehog signaling pathway, is dependent on the presence of primary cilia in melanoma cells (O’Reilly *et al*, 2013; Stecca *et al*, 2007). Correlating primary cilia formation with mutations in the *BRAF*, *PTEN*, and *NRAS* genes in melanoma may also clarify the value of using primary cilia to inform melanoma risk or survival.

## MATERIAL AND METHODS

### Cell lines

A375 (ATCC Cat. #CRL-1619), HEK293T (ATCC Cat. #CRL-11268), and YUMM1.7 (ATCC Cat. #CRL-3362) cells were cultured in DMEM, high glucose GlutaMAX™ with pyruvate supplement (Thermo Fisher Scientific Cat. #31966047) containing 10% FBS (Carlo Erba Reagents Cat. #FA30WS1810500) and 100 µg/ml penicillin and streptomycin (Thermo Fisher Scientific Cat. #15140122). Cells were maintained in a 37 °C incubator with humidified atmosphere of 5% CO2. All cell lines were regularly tested for mycoplasma infection using N GARDE Mycoplasma PCR Reagent Set (EuroClone Cat. #EMK090020).

### Ethics statement

All procedures involving animals for research purposes were approved by the Organismo preposto al benessere degli animali (O.P.B.A.) of the University of Padua and by the Italian Ministry of Health (Permit Numbers: PR n°983/2024, 532/2025).

### In vivo experiments

For xenografts experiments, 5×105 cells resuspended in 100 μL of equal amounts PBS:Matrigel (Growth Factor Reduced Matrigel, Corning Cat. #354230) were injected subcutaneously on the back of 8–10-week-old NODSCID or C57BL/6 mice using a syringe with a 26G needle. Tumor volume was measured three times a week with a digital caliper and calculated as Volume = (Length × Widthff^2^)/2. After the tumor reached a palpable size (300 mm^3^), mice were treated with control chow or chow containing combination therapy (4RF21 chow added with 150 mg/kg Dabrafenib and 6 mg/kg Trametinib, supplied from Mucedola S.R.L.).

### Lentivirus production

Specific human shRNAs targeting GOT1 and GOT2 were obtained from SigmaAldrich in a pLKO.1 backbone (shRNA IDs: TRCN000034784 for shGOT1, TRCN0000034827 for shGOT2). Lentiviruses encoding these shRNAs were produced in HEK293T packaging cells using packaging plasmids pMD2.G (VSVG envelope), pMDLg/pRRE (Gag/Pol), and pRSV (Rev) by PEI-mediated transfection in Opti-MEM medium (Thermo Fisher Scientific Cat. #51985026). Medium was replaced 16 hours post-transfection, and viral supernatants were harvested 48 hours later, collected by ultracentrifugation, and resuspended in 1% BSA in PBS. Aliquots were stored at -80°C. Melanoma cells were infected with respective lentivirus particles to induce knockdown (KD), the medium was replaced after 24 hours, and cells were collected or analysed according to the experiment. All procedures for lentiviral production and use were conducted in a biosafety level-2 (P2) environment.

### Cell proliferation/viability/toxicity assay

A number of 1500/3000 cells/well were seeded in 96-well plates. After adhesion, cells were treated according to the experiment. At the experimental time points, the medium was removed, and cells were washed twice with PBS. Then, 100 µl of 0.5% (w/v) Crystal Violet staining solution in 20% methanol was added to each well and incubated for 20 min at room temperature on a bench rocker with a frequency of 20 oscillations per minute. The plates were washed four times with tap water and allowed to dry. Pictures of each well were taken using a Leica MZ16F or Zeiss Axiovert 5 microscope, and crystal violet-stained cells were quantified using ImageJ.

### Cell Treatments

For the combination therapy, cells were treated with 250 nM Dabrafenib (Cat. #: HY-14660) and 10 nM Trametinib (Cat. #HY-10999). Media was changed 3 times a week. We selected drug-resistant cells by continuously exposing them to the combination therapy for over a month, or by selecting cells that proliferated again under drug pressure. For the combination therapy toxicity assays, cells were treated with increasing concentrations of the drugs, maintaining the same ratio between dabrafenib and trametinib as the combination therapy. For shh inhibition, cells were treated with increasing concentrations of GANT61 (Cat. #HY-13901). For glutamine starvation, cells were kept in DMEM without glutamine (Cat. #11960044) for the indicated time, or the same medium replenished with 200 mM glutamine (Cat. #25030081). For the rescue with dimethyl 2-oxoglutarate (DM-2OG) (Sigma, Cat. #349631), a final concentration of 2 mM was used.

### Colony formation Assay

Seed 1000 cells/well in 6-well plates in complete DMEM. After adhesion, cells were treated with combination therapy or control medium. The medium was replaced 3 times a week and the colony formation was assessed by staining with 0.5% (w/v) Crystal Violet staining solution (Sigma) in 20% methanol for 20 minutes at RT. The staining solution was removed, cells were washed twice with tap water and allowed to dry. Colonies were imaged and manually counted, considering colonies of at least 10 cells.

### Real-time qPCR

Cells were rinsed with ice-cold PBS and either QIAzol™ Reagent (Qiagen, Cat. #79306) was directly added to the cells. After sample collection, RNA isolation was performed according to the manufacturer’s instructions. RNA concentration and purity were determined using a NanoDrop spectrophotometer. RNA was reverse transcribed to cDNA using a HighCapacity cDNA Reverse Transcription Kit (Thermo Fisher Scientific Cat. #4368814). Quantitative PCR was carried out with 5x HOT FIREPol® EvaGreen® qPCR Supermix (Solis Cat. #08-36-00020) on QuantStudio™ 5 Real-Time PCR System (Thermo Fisher Scientific). Transcript levels were normalized to β-Actin and reported as relative to a reference sample. All primers used in q-PCR are listed below. QuantStudio™ Design & Analysis Software **Error! Hyperlink reference not valid.** was used for performing q-PCR data processing.

Human β-ACTIN FW: 5′-GATGGAGTTGAAGGTAGTTTCGT-3′; Human β-ACTIN RV: 5′-GCGGGAAATCGTGCGTAGCATT-3′; Human eIF4E3 FW 1: 5’-GCAAAGGGTGGCGTATGGAA-3’; Human eIf4E3 RV 1: 5’-CCCGATGGTTGCTAACAGC-3’; Human eIF4E3 FW 2: 5’-AGCAACCATCGGGGAACAG-3’; Human eIf4E3 RV 2: 5’-CCGAACACTGACACTAACTCCTA-3’; Human ATAT1 FW: 5’-GGCCCAGAATCTTTCCGCTC-3’; Human ATAT1 RV: 5’-GATGCAAAGTGGTTCTACCTCAT-3’; Human BBS4 FW: 5’-ATGCTGGGGAAGATCCACTTG-3’; Human BBS4 RV: 5’-TGGTAAATGCCGAGCTGTAAGTA-3’; Human DNAH5 FW: 5’-TCTGGAAGCATAGCGTCACTC-3’; Human DNAH5 RV: 5’-CCTCCACTTCGGTTTTGTTCA-3’; Human IFT22 FW: 5’-GCTCATGGAGTGGTGATCGTC-3’; Human IFT22 RV: 5’-TGTGGTGTGCAATTAGCATACA-3’; Human PKD2 FW: 5’-CTCTGGGGAACAAGACTCATGG-3’; Human PKD2 RV: 5’-TCATCATGCCGTAGGTCAAGA-3’; Human TMEM237 FW: 5’-AGACTTCCTCCACCCAAAAGA-3’; Human TMEM237 RV: 5’-CTGGCTCCGCATCAATACCAT-3’; Human ITGAV FW: 5’-GCTCAGAGGGAGAGGAGCCTG-3’; Human ITGAV RV: 5’-AGTAACCCTGTCCTGCTGCC-3’; Human ITGA2 FW: 5’-TGTGGTGAGGACGGACTTTG-3’; Human ITGA2 RV: 5’-CATCAACCGGCAGGGAGAAT-3’;

### Western Blotting

Cells were rinsed with ice-cold PBS and collected from culture plates by scraping. Cells were lysed in ice-cold homemade RIPA Lysis Buffer (20 mM TrisHCl, pH 7.4, 200 mM NaCl2, 2% sodium deoxycholate, 0.2% SDS, 2% NP40) supplemented with 1x halt protease inhibitor cocktail (Roche Cat. #04693116001) and phosphatase inhibitor (Sigma Cat. #4906845001) for 20 min, and soluble lysate fractions were clarified by centrifugation at 20,000 × g for 10 min at 4 °C. In case of puromycin-incorporation assays, cells were treated with 50 μg/ml puromycin (Sigma Cat. #P8833) for 10 minutes at 37°C prior to collection. Protein concentrations were determined with Pierce™ BCA Protein Assay Kit (Thermo ScientificTM, Cat# 23225). 10 μg of protein per lane was loaded in SDSPAGE (4-12% pre-cast gels Invitrogen) and transferred to nitrocellulose membrane. After blocking with 5% milk in TBS-T buffer for 1h, the membranes were incubated with the primary antibodies overnight at 4°C in 1% BSA. Following the incubation, all membranes were washed three times in TBS-T buffer prior to incubation with the appropriate horseradish peroxidase (HRP)- conjugated secondary antibodies (IgG) (1:10 000). The bands were visualized using Immobilon Forte Western HRP substrate and acquired using ChemiDoc™ Imaging System. The following primary antibodies were used: anti-human VINCULIN (1:5000, Sigma Cat. #V9131, RRID:AB_477629); Puromycin (1:10,000, Merck Cat. #MABE343, RRID:AB_2566826); Phospho-S6 Ribosomal Protein (Ser240/244) (D68F8) (1:1000, Cell Signalling Cat. #5364, RRID:AB_10694233); S6

Ribosomal Protein (5G10) (1:1000, Cell Signalling Cat. #2217, RRID:AB_331355); Phospho-p70 S6 Kinase (Thr389) (1:1000, Cell Signalling Cat. #9205, RRID:AB_330944); p70 S6 Kinase (49D7) (1:1000, Cell Signalling Cat. #2708, RRID:AB_390722); Rabbit polyclonal anti-ASNS (1:1000, Cell Signaling Technology,Cat. #20843); Rabbit polyclonal anti-Glutaminase (1:1000, Abcam Cat.#ab93434; RRID: AB_10561964); Rabbit monoclonal anti- 68 GOT1 (clone E4A40) (1:1000, Cell Signaling Technology, Cat. #34423); Rabbit polyclonal anti-GOT2 (1:1000, Thermo Fisher Scientific, Cat. #77990; RRID: AB_2735868); Rabbit polyclonal anti-p44/42 MAP kinase (phosphorylated Erk1/2)(1:1000, Cell Signaling Technology, Cat.#9101; RRID: AB_331646); Mouse monoclonal anti-Erk1/2 (clone C-9) (1:1000, Santa Cruz Biotechnology, Cat.#sc514302; RRID: AB_2571739); Rabbit Glutamine synthetase antibody (1:1000, GeneTex, Cat. #GTX109121); Rabbit IFT88 Polyclonal antibody (1:1000, ProteinTech, Cat. #13967-1-AP); eIF4E Rabbit mAb (1:1000, ABClonal, Cat. #A25608); Rabbit EIF4E3 Polyclonal antibody (1:1000, ProteinTech, Cat #17282-1-AP); HIF1 alpha rabbit polyclonal antibody (1:1000, GeneTex, Cat.# GTX127309); Phospho-AMPK alpha (Thr172) (40H9) Rabbit Monoclonal Antibody (1:1000, Cell Signaling, Cat.# 2535); RRM2 (E7Y9J) Rabbit Monoclonal Antibody (1:1000, Cell Signaling, Cat.# 65939); Monoclonal Anti-Cyclin A mouse antibody (1:1000, Sigma-Aldrich, Cat.# C4710); ATF-4 (D4B8) Rabbit Monoclonal Antibody (1:1000, Cell Signaling, Cat.# 11815); RPL23 Polyclonal Antibody (1:1000, Invitrogen, Cat.# PA5-110318); Ribosomal Protein L26 Antibody (1:1000, Cell Signaling, Cat.# 2065); Ribosomal Protein S3 (D50G7) Rabbit Monoclonal Antibody (1:1000, Cell Signaling, Cat.# 9538); AMPK alpha rabbit Antibody (1:1000, Cell Signaling, Cat.# 2532).

### RNA and Ribo-seq sequencing A375

Matched bulk RNA sequencing data (RNA-seq) and ribosome profiling (Ribo-seq) libraries were processed using the same human reference genome and gene annotation (GENCODE human v42). Reads from both RNA-seq and Ribo-seq libraries were aligned to the reference genome using STAR Aligner. Gene-level read quantification was performed using the htseq-count module of HTSeq with the --mode=intersection-strict option. For RNA-seq, reads overlapping annotated exons were quantified using --type=exon, whereas Ribo-seq reads were quantified over coding sequences using --type=CDS. The resulting gene-level count matrices were used for downstream differential expression and translation efficiency analyses.

### Differential expression analysis of A375

Gene-level read counts from RNA-seq data in A375 cells were processed using DESeq2 v1.50.2(Love *et al*, 2014). Genes with low expression were removed by retaining only those with a minimum count threshold of ≥5 reads in at least 3 samples. Differential expression analyses were performed using a single-factor design (∼ condition). log2(fold changes) were shrunk using apeglm v1.32.0 (Zhu *et al*, 2019). Genes were considered significantly up and down-regulated for an adjusted p-value < 0.1 and a |log2(fold change)| > 0.5.

### Differential expression analysis of GSE77940

Gene expression quantification data for matched pre- and post-treatment melanoma samples were retrieved from dataset GSE77940 (Wagle *et al*., 2014). Non-expressed and low-expressed genes were filtered by retaining only those with an expression level exceeding 1 fragments per kilobase of transcript per million (FPKM) in at least 6 of the 12 total samples. Expression values subsequently transformed using a log2(FPKM+1) scale. Differential expression was performed using the limma v3.68.4 (Ritchie *et al*, 2015) (package with a paired linear design (∼ patient + treatment). Linear models were fitted for all filtered genes using lmFit, followed by empirical Bayes variance moderation with eBayes.

### Gene Set Enrichment Analysis (GSEA) for GSE77940 and A375

Pre-ranked GSEA for GSE77940 was evaluated applying the clusterProfiler v4.18.4 package (Yu *et al*, 2012) using the moderated t-statistic. Human gene set annotation collections were retrieved via the msigdbr package v26.1.0 (Dolgalev, I. (2022). msigdbr: MSigDB Gene Sets for Multiple Organisms in a Tidy Format. R package version 7.5.1. https://CRAN.R-project.org/package=msigdbr), covering Reactome pathways (Milacic *et al*, 2024) or Gene Ontology Biological Processes (GO-BP) (Ashburner *et al*, 2000) and KEGG pathways (Kanehisa *et al*, 2023). GSEA was executed independently across each gene set collection using the Benjamini-Hochberg method to control the false discovery rate. For pre-ranked GSEA for A375, all detected genes were ranked in descending order based on their DESeq2 Wald test statistic. GSEA was executed across the databases of Gene Ontology Biological Processes (GO-BP) (Ashburner *et al*., 2000)(GO-BP) using gseGO, and KEGG pathways using gseKEGG applying the Benjamini-Hochberg procedure for FDR control. Significant and top-ranking enriched pathways were visualized for all datasets using dot plots stratified by the sign of the Normalized Enrichment Score (NES).

### Over-representation analysis (ORA) for A375

ORA was performed using over-representation analysis (ORA) via clusterProfiler v4.18.4 (Yu *et al*., 2012). Significantly up- and down-regulated genes (adjusted p-value < 0.1, |log_2_(fold change)| > 0.5) were tested against Gene Ontology Biological Processes (GO-BP) (Milacic *et al*., 2024) using enrichGO. Significantly enriched GO-BP terms derived from ORA were visualized using dot plots. The top 20 most significant categories (ranked by adjusted p-value, Benjamini-Hochberg procedure for FDR control) were selected for each subset of up- and down-regulated genes.

### Ribosome occupancy

P-site coordinates were assigned from aligned Ribo-seq reads aligned to the transcriptome using riboWaltz (Lauria *et al*, 2018) based on footprint length-specific offset distances. Structural annotations of the transcripts were extracted to map positional P-site coverage across the genes. Positional P-site counts were first normalized to footprint library size as Reads Per Million mapped reads (RPM). For coding regions, nucleotide-level RPMs were summed across non-overlapping 3-nt frames to yield codon-resolution profiles, while 5′ UTR profiles were evaluated at nucleotide resolution relative to the canonical AUG start codon. To decouple ribosome density from mRNA expression, positional footprint RPMs were divided by matched gene-level RNA-seq abundance (Counts Per Million, CPM) to generate normalized occupancy scores (P-site RPM/RNA CPM). Profiles were smoothed using a moving-average filter and summarized across biological replicates as mean ±SEM for conditions and visualized with ggplot2 (Wickham, Hadley, and Carson Sievert. ggplot2: elegant graphics for data analysis. Vol. 10. New York: springer, 2009).

### Codon-resolved ribosome occupancy

Ribosome footprints were mapped to the transcriptome (GENCODE v42), and P-site offsets were determined per read length with riboWaltz. For codon-level analysis, only P-sites located within the annotated coding sequence (CDS) and in frame 0 were retained, restricted to the transcript with the longest CDS of each gene. The codon at each P-site was read from the transcript sequence, and stop codons were excluded. For each sample, the RNA CPM of the corresponding gene was assigned to the transcript. Transcripts with CPM < 1 in the respective sample were excluded. P-site counts of each codon in each transcript were divided by (CPM + 0.01), where the pseudocount avoids division by zero, to obtain ribosome occupancy per unit of mRNA. These values were summed over all transcripts for each codon. The resulting codon values were converted to fractions of the total, divided by the frequency of the respective codon in the CDSs of the analysed transcripts, and expressed relative to the mean across the 61 sense codons, so that a value of 1 indicates uniform occupancy across codons. Values were calculated for each sample and averaged across the three biological replicates of each condition.

### Generation of IFT88 knockout clones

IFT88 knockout (KO) clones were generated in A375 cells by CRISPR/Cas9- mediated genome editing. A single guide RNA (gRNA) targeting IFT88 (5’-GAACTTGGTCTAATGCCATT-3’) was cloned into the pX459 plasmid (pSpCas9(BB)-2A-Puro V2.0; Addgene #62988), as described before(Ran *et al*, 2013). A375 cells were transfected with the resulting construct using Lipofectamine 2000 70 (Invitrogen, cat #11668027), following the manufacturer’s protocol. At 48 h post-transfection, transfected cells were selected with puromycin (1 µg/mL) for 3 days to enrich for cells carrying the construct. Following selection, single cells were seeded into 96-well plates by limiting dilution to establish clonal populations. Once individual clones had expanded sufficiently, IFT88 protein expression was validated by western blot.

### Polysome profiling analysis

For polysome profiling, A375N and A375DR cells were seeded in triplicate in 15-cm dishes. When cultures reached ∼80% confluence, cells were treated with cycloheximide (CHX; 100 µg/mL) for 10 min. Cells were then gently washed with ice-cold PBS containing 10 µg/mL CHX and lysed in buffer composed of 30 mM Tris-HCl (pH 7.5), 100 mM NaCl, 30 mM MgCl_2_, 0.1% NP-40, 100 µg/mL CHX, and 30 U/mL RNasin. Lysates were clarified by centrifugation at 13,000 × g for 10 min at 4 °C. Cytoplasmic extracts containing equal amounts of RNA were loaded onto 15–50% (w/v) buffered sucrose gradients (20 mM Tris-HCl, 140 mM KCl, 10 mM MgCl_2_, 1 mM DTT, 100 µg/mL CHX; pH 7.4) prepared in 14 mm × 89 mm thin-wall Ultra-Clear tubes (Beckman #344059) using a Biocomp Gradient Master. Gradients were centrifuged at 35,000 rpm for 120 min at 4 °C in a SW-41Ti rotor (Beckman) with maximum acceleration and no brake on a Beckman Optima L-80K ultracentrifuge. Gradients were fractionated using a Biocomp piston fractionator equipped with a TRIAX flow cell, while continuously recording absorbance at 260 nm.

### Cell cycle analysis

Cell cycle was analyzed by flow cytometry using propidium iodide staining. After the indicated treatments, cells were detached by trypsinization (EuroClone, Cat. #ECB3052). The cell pellets were washed with PBS, then resuspended in 300 µL of PBS. Propidium iodide (Sigma, cat. #P4864) was added at a final concentration of 10 µg/mL prior to analysis. Flow cytometry was performed using the BD FACSCanto™ II Cell Analyzer (BD Biosciences).

### OCR and ECAR analyses

Oxygen consumption rate (OCR) and ECAR (Extracellular Acidification Rate) was measured using a Seahorse XF24 Extracellular Flux Analyzer (Agilent Technologies) with the XF Cell Mito Stress Test Kit. Cells were seeded at 3 × 10^4^ cells/well in XF24 cell culture microplates 24 h before the assay. Four wells in the standard background positions (A1, B4, C3, D6) were left without cells for background correction. The sensor cartridge was hydrated overnight with XF Calibrant at 37 °C in a non-CO_2_ incubator.

On the day of the assay, the culture medium was replaced with HEPES-buffered XF DMEM assay medium (pH 7.4), and cells were equilibrated at 37 °C in a non-CO_4_ incubator before measurement. Following baseline measurements, oligomycin (1.5 µM), FCCP (1 µM) and rotenone/antimycin A (0.5 µM) were sequentially injected using the instrument’s standard pre-set Mito Stress Test measurement cycles.

After the assay, nuclei of live cells were stained and imaged with an Operetta high-content imaging system, and cell number per well was determined with Harmony software (Revvity). OCR data were normalized to cell number and expressed as pmol O_2_/min/10⁴ cells. Normalization was performed with Agilent Seahorse Analytics (online software).

### Immunofluorescence and cilia quantification

Isolated tumors were fixed in 4% PFA overnight. Then they were incubated in 20% sucrose for 2 days. Tumor cryosections (10 µm-thick) were blocked in PBS containing 5% goat serum (Merk Millipore Cat. #S26), 1% BSA and 0.3 M glycine for 30 min. Sections were then incubated with Acetyl-alpha-Tubulin (Lys40) (D20G3) Rabbit Monoclonal Antibody (1:300, Cell Signalling Technology, Cat. #5335) and mouse ARL13B (C-5) antibody (1:150, Santa Cruz, Cat. #sc-515784) overnight at 4 °C. After three washes with PBS, sections were incubated for 1 h at RT with the corresponding Alexa Fluor conjugated secondary antibodies (5 µg/ml, Thermo Fisher Scientific) and 300 nM DAPI (Thermo Fisher Scientific Cat. #D1306) for nuclear staining. Sections were mounted and then analyzed using Leica SP8 DLS microscopy. For cultured cell immunofluorescence, cells were seeded on glass coverslips placed in a 24-wells plate. Cells were washed three times with PBS and fixed with 4% paraformaldehyde for 15 min at RT. The fixed cells were then permeabilized with 0.2% Triton X-100 in 1× PBS for 10 min at RT and blocked with 3% goat serum + 3% BSA + 1.5 mg/ml glycine in PBS for 1 h at room temperature. Cells were incubated overnight at +4 °C with the primary antibody. After three washes with PBS, cells were incubated for 1 h at room temperature with the corresponding Alexa Fluor conjugated secondary antibodies (0.75– 2 µg/ml, Thermo Fisher Scientific) in 0.2% BSA/PBS and 300 nM DAPI (Thermo Fisher Scientific) for nuclear staining. After three additional washes with PBS, coverslips were mounted with Mowiol® 4-88 (Sigma) and then visualized using Leica SP8 DLS microscopy. Images were processed using Leica LASX software. Primary cilia were quantified with the ACDC software(Lauring *et al*, 2019).

### Targeted metabolomics

Metabolites were extracted by lysing cell pellets in 250 μL of ice-cold methanol/acetonitrile/water (55:25:20, v/v/v) containing 1 ng/μL of [U-13C6]-glucose (Sigma-Aldrich, cat. no. 389374), [U-13C5]-glutamine (Sigma-Aldrich, cat. #605166), and 0.2 ng/μL of [U-13C2, 15N]-glutathione (GSH) (SigmaAldrich, cat. #683620) as internal standards. Samples were centrifuged at 15,000 × g for 15 min at 4 °C, and the supernatants were collected and dried under a nitrogen stream at 40 °C. Dried extracts were resuspended in 120 μL of methanol/water (70:30, v/v) prior to analysis. For amino acid quantification, 10 μL of each extract were derivatized with 50 μL of a 1:1:1 (v/v/v) mixture of H_2_O, ethanol, and pyridine containing 5% phenylisothiocyanate (PITC). Following incubation for 20 min at room temperature, samples were dried under nitrogen and reconstituted in 100 μL of 5 mM ammonium acetate/methanol-water (1:1, v/v). Targeted metabolite quantification was performed by liquid chromatography tandem mass spectrometry (LC-MS/MS). Amino acids were separated using a Biocrates C18 column maintained at 50 °C, acyl-carnitines using a ZORBAX StableBond CN column (2.1 × 150 mm, 5 μm; Agilent), and metabolites, cofactors, and nucleotides using a Luna CN column (50 × 4.6 mm, 5 μm; Phenomenex) maintained at 35 °C. Amino acids and acyl-carnitines were analyzed in positive electrospray ionization (ESI+) mode using mobile phase A (0.2% formic acid in water) and mobile phase B (0.2% formic acid in acetonitrile) with a gradient of 100% A (0 min), 5% A (5.5 min), and 100% A (7 min). Flow rates were 500 μL/min for amino acids and 300 μL/min for acyl-carnitines. All other metabolites were analyzed in negative ESI mode using water as mobile phase A and 2 mM ammonium acetate in methanol as mobile phase B under isocratic conditions (50% B) at a flow rate of 600 μL/min. A total of 100 metabolites were identified using multiple reaction monitoring (MRM) transitions previously validated with pure standards during method development. Data were acquired on an API 3500 triple quadrupole mass spectrometer (AB Sciex) coupled to an HPLC system. Peak integration and quantification were performed using MultiQuant™ software (version 3.0.2). Raw peak areas were median-normalized, log-transformed, and Pareto-scaled prior to statistical analysis using MetaboAnalyst 6.0. To compare relative abundance of single metabolites, no log transformation and pareto-scaling was applied after median-normalization(Pang *et al*, 2024).

### Statistical Analysis

Statistical analyses were performed using GraphPad Prism (version 10), unless otherwise specified. Data are presented as mean ± standard deviation (SD), except for in vivo tumor growth curves, which are shown as mean ± standard error of the mean (SEM). Comparisons between two groups were performed using an unpaired two-tailed t-test. Comparisons among more than two groups were performed using one-way ANOVA, or two-way ANOVA when two factors were considered, followed by Tukey’s or Šídák’s multiple comparisons test, as indicated in the figure legends. Growth curves were fitted with an exponential (Malthusian) growth model by least squares regression, and doubling times were compared using the extra sum-of-squares F test. Dose-response curves were fitted using the “[Inhibitor] vs. response (three parameters)” model by least squares regression, and IC50 values were compared using the extra sum-of-squares F test. Metabolomic data were analyzed as described in the “Metabolomics" section. In figures, asterisks indicate statistical significance (*p < 0.05, **p < 0.01, ***p < 0.001, ****p < 0.0001), where p refers to the unadjusted p-value for t-tests and F tests, and p-adj for ANOVA post hoc tests and for the false discovery rate (FDR)-adjusted p-values in the metabolomic analysis (*p adj < 0.05, **p adj < 0.01, ***p adj < 0.001, ****p adj < 0.0001). ns, not significant.

## Author contribution

M.M.S. and P.L. conceived and designed the study; M.M.S. supervised and wrote the manuscript. P.L. performed the in vivo and in vitro work on melanoma cells. R.C. performed in vivo YUMM experiments; C.T. performed bioinformatic analyses. R.O. performed polysome profiling experiments; S.P. and N.C. performed and interpreted MS metabolic analysis. All authors read and approved the final paper.

