## Supplementary figures and images for "THERAPY RESISTANCE IN MELANOMA DRIVES CILIOGENESIS AND GLUTAMINE/ASPARTATE METABOLISM DEPENDENCY"

### Supplementary Figure

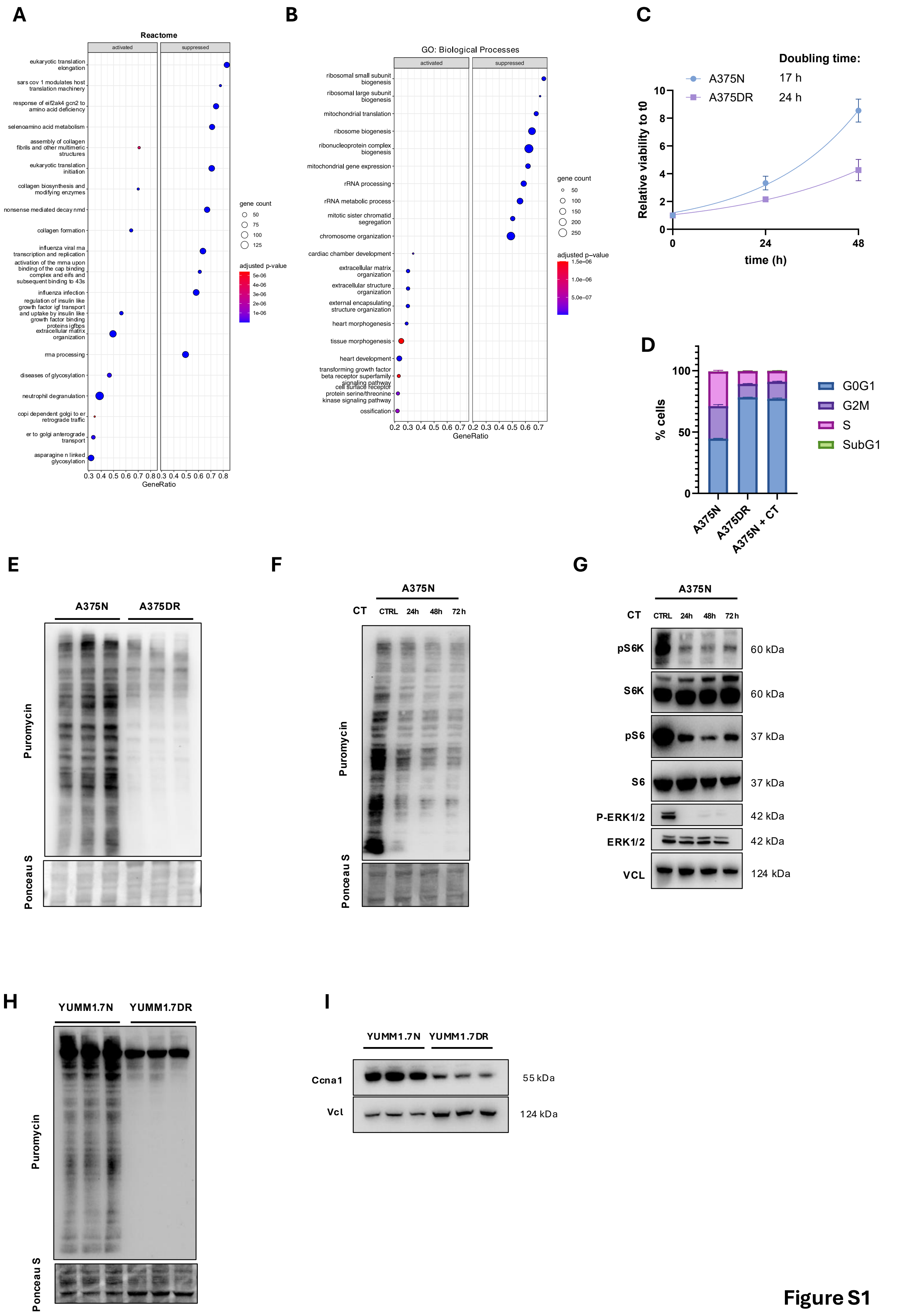

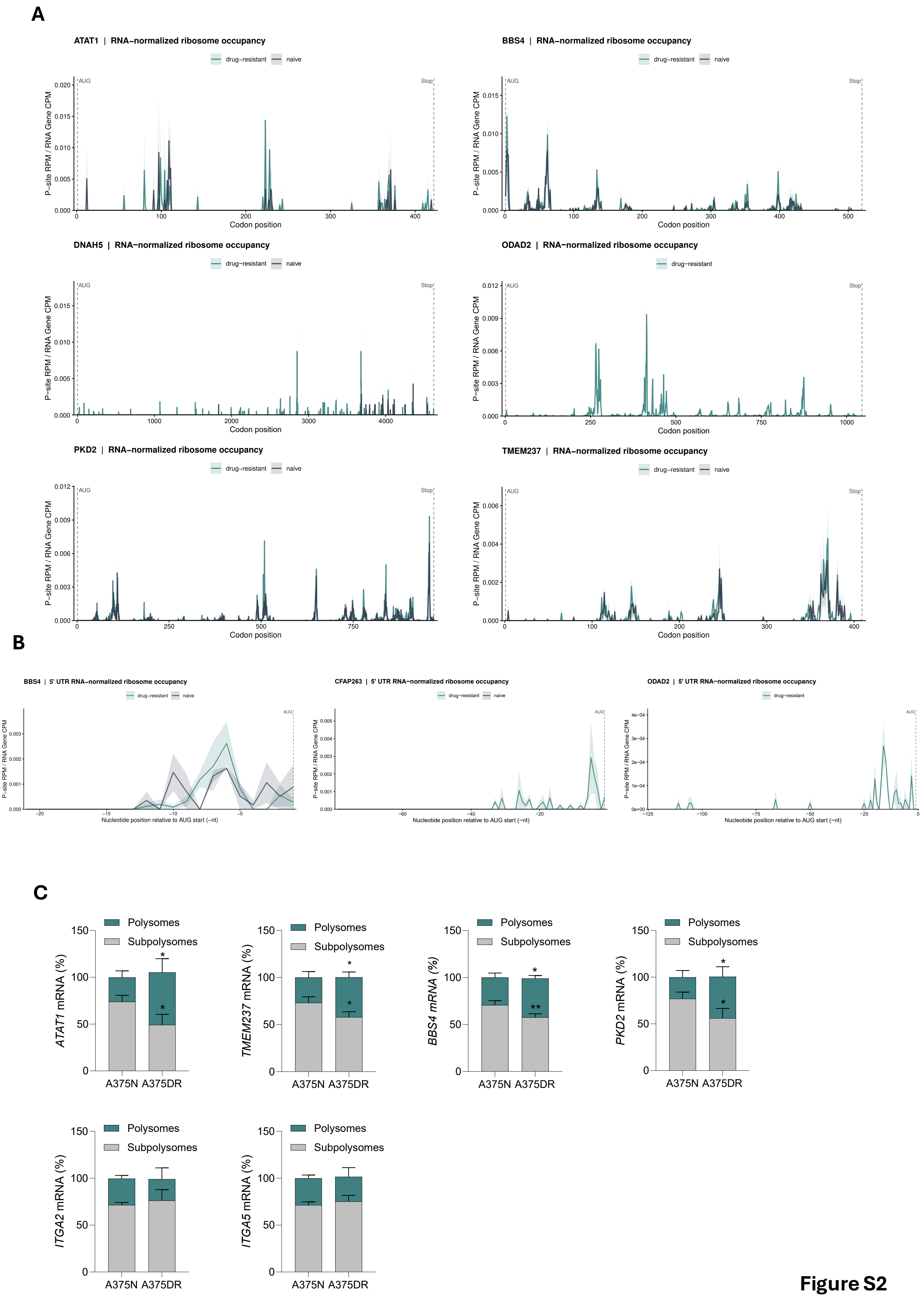

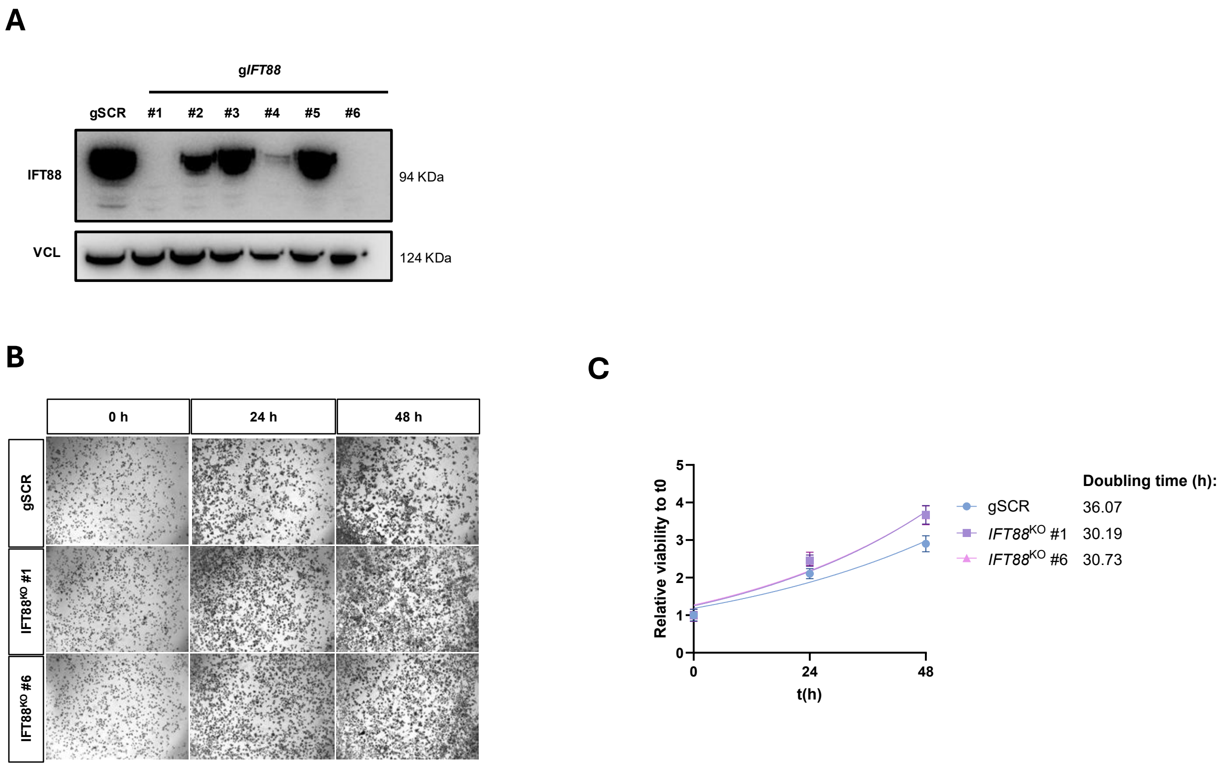

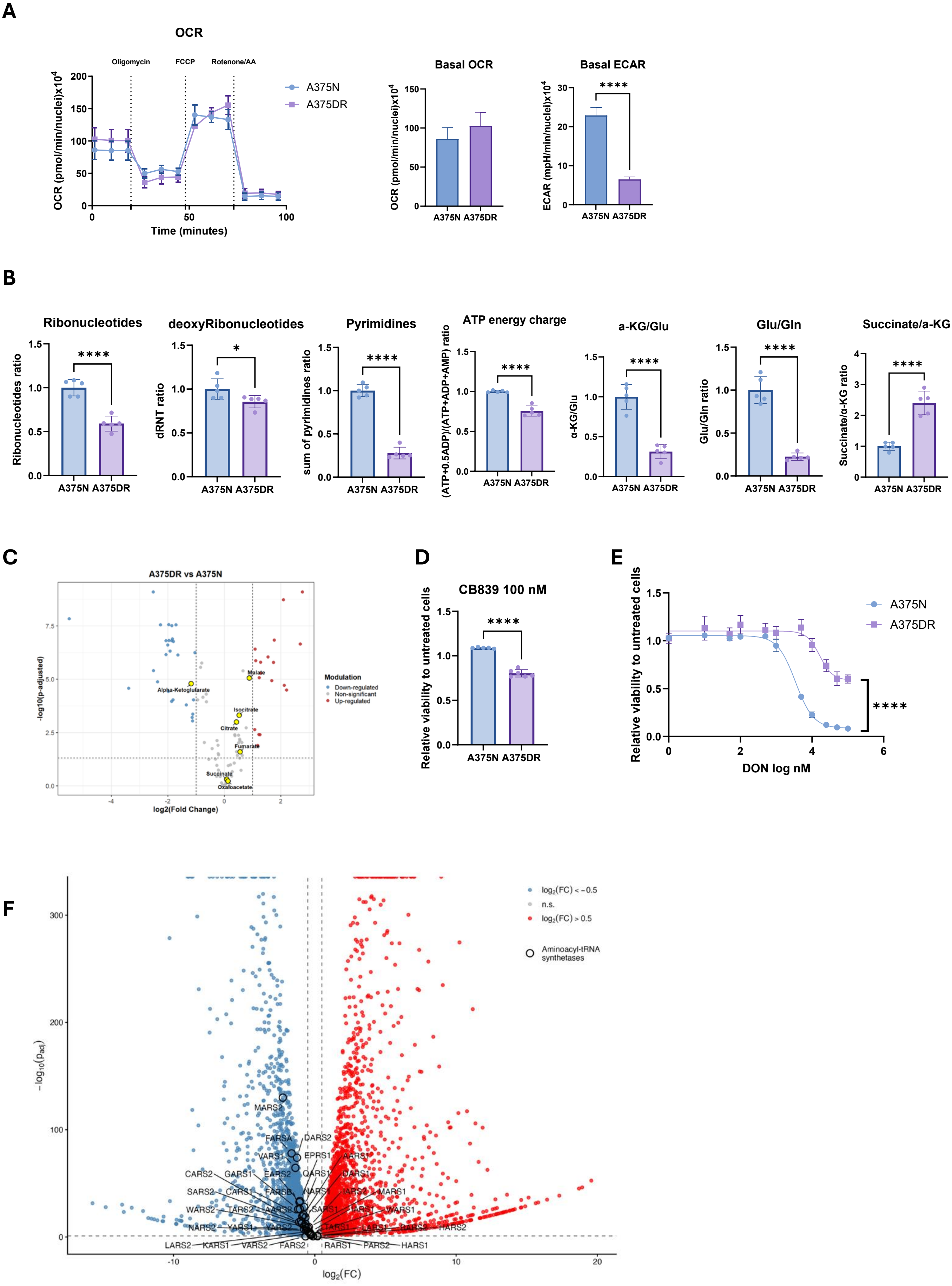

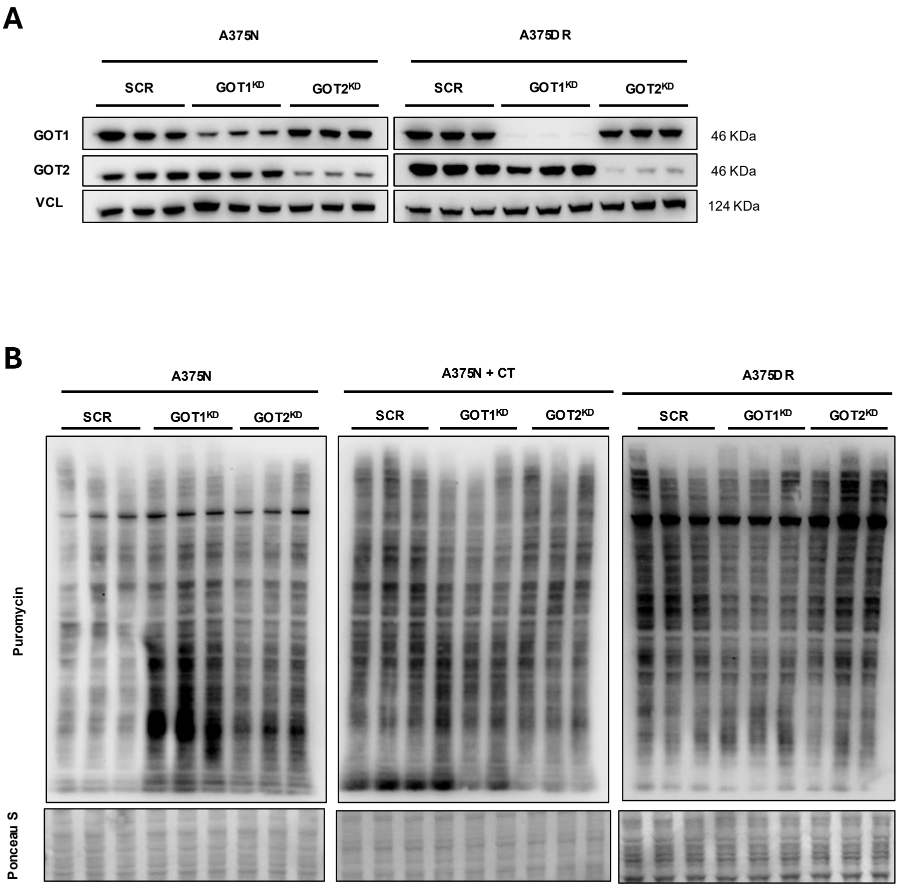

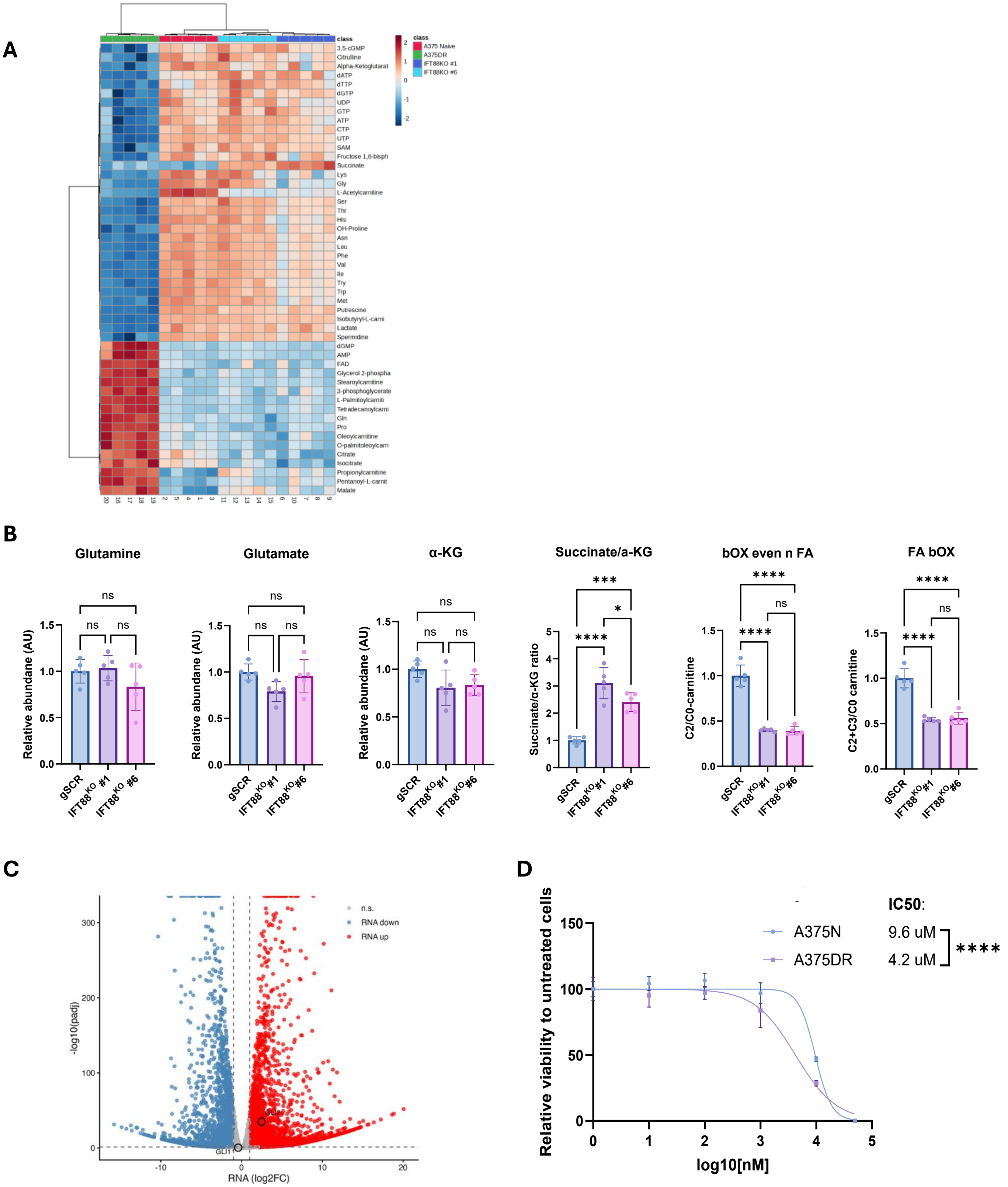
